# Mediator Kinase Inhibition Impedes Transcriptional Plasticity and Prevents Resistance to ERK/MAPK-Targeted Therapy in *KRAS*-Mutant Cancers

**DOI:** 10.1101/2022.09.17.508384

**Authors:** Daniel P. Nussbaum, Colin A. Martz, Andrew M. Waters, Alejandro Barrera, Justine C. Rutter, Christian G. Cerda-Smith, Amy E. Stewart, Chao Wu, Merve Cakir, Cecilia B. Levandowski, David E. Kantrowitz, Shannon J McCall, Mariaelena Pierobon, Emanuel F. Petricoin, J. Joshua Smith, Timothy E. Reddy, Channing J. Der, Dylan J. Taatjes, Kris C. Wood

**Author notes:** First author (Daniel P. Nussbaum).

## Abstract

Acquired resistance remains a major challenge for therapies targeting oncogene activated pathways. *KRAS* is the most frequently mutated oncogene in human cancers, yet strategies targeting its downstream signaling kinases have failed to produce durable treatment responses. Here, we developed multiple models of acquired resistance to dual-mechanism ERK/MAPK inhibitors across *KRAS*-mutant pancreatic, colorectal, and lung cancers, and then probed the long-term events enabling survival against this novel class of drugs. These studies revealed that resistance emerges secondary to large-scale transcriptional adaptations that are diverse and tumor-specific. Transcriptional reprogramming extends beyond the well-established early response, and instead represents a dynamic, evolved population-level process that is refined to attain a stably resistant phenotype. Mechanistic and translational studies reveal that resistance to dual-mechanism ERK/MAPK inhibition is broadly susceptible to manipulation of the epigenetic machinery, and that Mediator kinase, in particular, can be co-targeted at a bottleneck point to prevent diverse, tumor-specific resistance programs.

## Introduction

Strategies to durably inhibit the RAF-MEK-ERK mitogen-activated protein kinase (MAPK) signaling network have the potential for broad use in cancers with *RAS* or *RAF* family activating mutations, amplifications in downstream kinases, and general dependence on MAPK signaling without genomically conspicuous pathway alterations. While pharmacologic inhibition of most *RAS*-mutant isoforms has to date proven elusive (*1, 2*), therapies targeting RAF, MEK, and ERK have demonstrated strong preclinical efficacy (*3, 4*), and in some cases, impressive activity in clinical trials (*5, 6*). Nonetheless, resistance to first generation ERK/MAPK inhibitors has been problematic, largely secondary to mutational and non-mutational pathway reactivation (*4, 7*), and strategies that dually inhibit ERK/MAPK or parallel signaling pathways have typically proven transient or clinically toxic in patients (*8–10*). Thus, ongoing efforts have focused on the development of strategies with the potential to potently, sustainably, and tolerably block ERK/MAPK signaling.

Novel classes of dual-mechanism RAF, MEK, and ERK inhibitors have recently been developed to more effectively inhibit ERK/MAPK signaling, predominantly by preventing known adaptations that lead to pathway reactivation (*11–16*). For example, the second generation pan-RAF inhibitor LY3009120 prevents paradoxical reactivation by RAF dimers (*11*), the MEK inhibitor GDC-0623 prevents feedback phosphorylation by wild-type RAF (*12*), and the allosteric and ATP-competitive ERK inhibitor SCH772984 enables catalytic blockade that overcomes phosphorylation by MEK, maintaining signaling abrogation despite negative feedback activation (*13*). Nevertheless, acquired resistance to ERK/MAPK inhibition with these and related agents has been increasingly reported (*17–19*), including in work from our laboratory which described evolved resistance to these kinase inhibitors across diverse cell line and animal models (*20*). This suggests that signaling events beyond traditional pathway reactivation support the development of resistance to more sustained ERK/MAPK suppression.

Recent studies have described dynamic enhancer remodeling in response to kinase inhibition, suggesting that acute chromatin events buttress diverse transcriptional escape programs, providing a window of opportunity to broadly constrain acquired drug resistance (*21, 22*). Rusan et al. demonstrated that the CDK7/12 inhibitor THZ1 can be leveraged to enhance cell death and suppress the emergence of resistance to inhibitors targeting oncogenic kinases (*21*). Similarly, Zawistowski et al. showed that BET bromodomain inhibition with JQ1 or I-BET151 can prevent resistance to the MEK inhibitor trametinib in MAPK-dependent triple negative breast cancer models by interfering with enhancer remodeling (*22*). These findings build off prior work demonstrating the ubiquity of transcriptional changes in drug resistance (*23–25*), and the concept of epigenetic remodeling to support adaptive, stable transcriptional programs (*26, 27*). However, mechanistic and translational questions remain regarding the combination of targeted kinase inhibitors with epigenetic manipulation. These pertain to 1) the nodes in an oncogenic pathway most susceptible to these combination strategies, 2) the specific DNA-interacting proteins most targetable in a given genomic and therapeutic context, and 3) the kinetics of acute versus stable epigenetic and transcriptional changes that facilitate survival and ultimately enable the terminally resistant state.

Here, we developed multiple models of acquired resistance to second generation, dual-mechanism ERK/MAPK inhibitors to characterize transcriptionally-mediated resistance in diverse *KRAS*-mutant cancer types. This oncogenic context was selected given the frequency of *KRAS* mutations in the most prevalent and lethal solid tumors (*28–30*), the refractory nature of *KRAS*-mutant tumors to single-agent ERK/MAPK inhibition (*7–10*), and the more recent observation that newer ERK/MAPK inhibitors are nonetheless susceptible to evolved resistance in a manner suggestive of transcriptional bypass programs (*17–20*). Counter to earlier models, we found that population-level transcriptional reprogramming extends beyond the well-established early response, and instead represents a dynamic, evolved process as cell populations refine their expression changes to attain a stably resistant phenotype. We delineate the previously hypothesized concept that the intrinsic early transcriptional response can be targeted as a “bottleneck” event and show how this period is broadly vulnerable to pharmacologic perturbation of chromatin-level events in order to block diverse, tumor-specific transcriptional resistance programs. Mechanistic and translational studies reveal that Mediator kinase inhibition antagonizes the initial early response to ERK/MAPK inhibition—which is highly enriched for genes within fundamental anabolic cellular processes—resulting in paralysis of the further population-level transcriptional events necessary for stable resistance. These findings demonstrate that co-targeting of Mediator kinase represents a novel, well-tolerated strategy for preventing resistance to sustained ERK/MAPK inhibition, and furthers our understanding of the kinetics and plasticity underlying drug response.

## Results

### A revised model of long-term resistance to sustained ERK/MAPK inhibition

We have previously assessed long-term ERK/MAPK inhibition across various models of *KRAS*-mutant cancers, and have consistently observed only transient sensitivity, followed by acquired resistance that develops over the course of several weeks (*20*) (Figure S1A, B). Novel inhibitors of RAF and MEK, despite their proposed mechanisms, nonetheless appear vulnerable to traditional pathway reactivation (*4, 31, 32*). Alternatively, while dual-mechanism ERK inhibition also initially activates well-described endogenous feedback events, compensatory feedback activation of MEK and ERK ultimately abates on the timescale at which stable resistance develops (Figure S1C), suggesting that distinct events might permit resistance to more sustained ERK/MAPK inhibition.

To confirm these findings in a more translationally relevant system, we utilized a well-credentialed, genetically engineered mouse model (GEMM)-derived *Kras*^G12D^/*Tp53*^-/-^ orthotopic, syngeneic mouse model of pancreatic cancer to assess resistance to dual-mechanism ERK inhibition (*33*) (Figures 1A-C). Like cell line models, murine tumors demonstrated initial drug sensitivity, yet developed resistance—as evidenced by progressive growth on treatment—within two weeks of treatment initiation (Figure 1A). This period of initial drug sensitivity was characterized by ineffective pathway reactivation, as evidenced by increased MEK phosphorylation at residues S217 and S221 without commensurate recovery of cell proliferation. Conversely, as stable resistance was achieved, flux through ERK/MAPK was relinquished (i.e. decreased phosphorylation of MEK and ERK), coupled by cell cycle reentry and tumor proliferation (Figure 1B,C). We observed similar findings in additional *in vitro* models of *KRAS*-mutant pancreatic and lung cancers (Figure S1D). Notably, these signaling events differ from those induced by first generation, single-mechanism ERK inhibitors, in which resistance is characterized by hyperphosphorylation of ERK at residues T202 and Y204 (Figure S1E). These findings suggest that while sustained ERK inhibition is susceptible to adaptive resistance, these programs rely on signaling events distinct from traditional ERK/MAPK pathway reactivation. Moreover, these observations provoke questions regarding the long-term role of MAPK signaling feedback events, which to date have been described on the scale of only hours to days in the context of acquired resistance (*4, 7*).

**Fig. 1.**
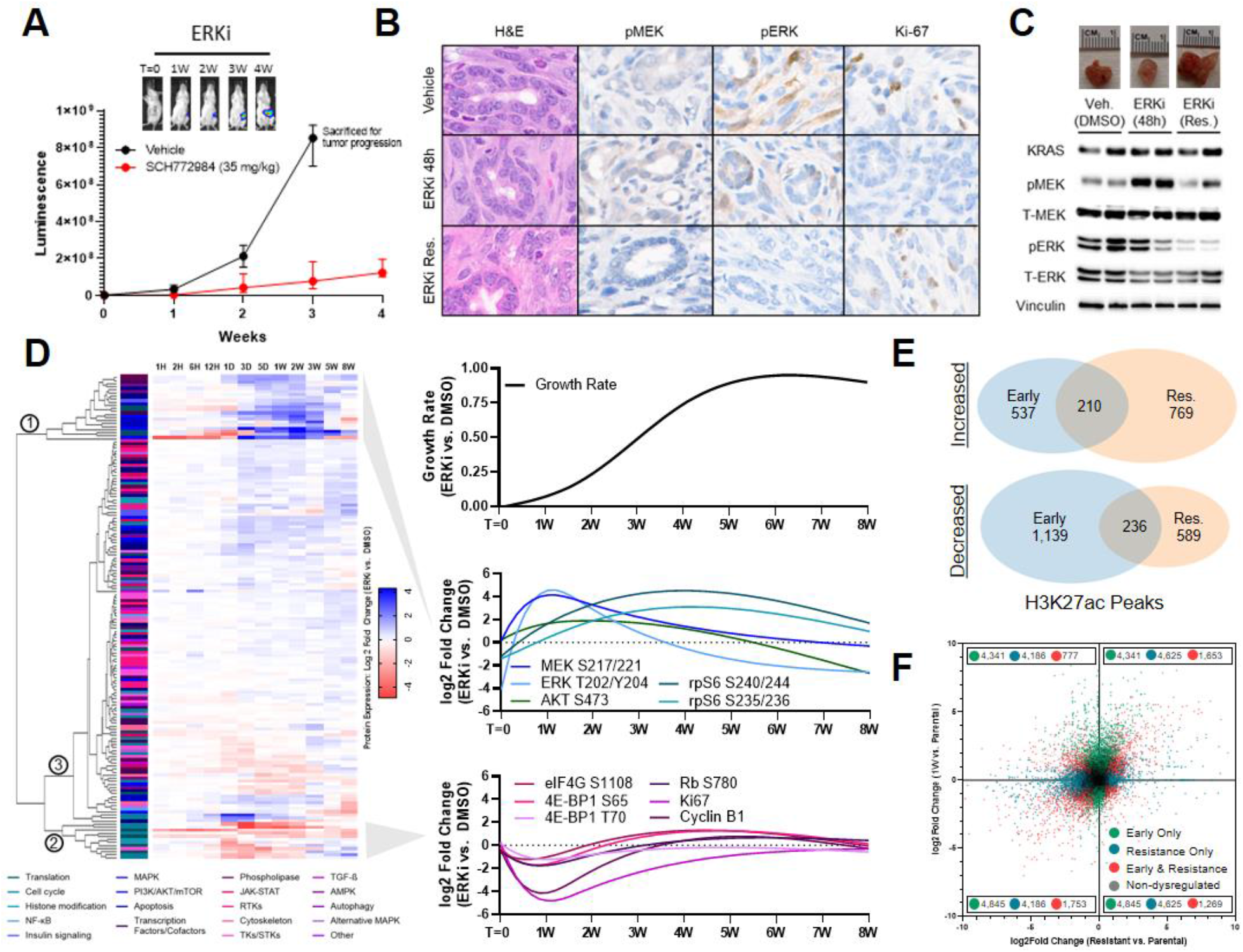
Resistance to ERK/MAPK inhibition is supported by long-term epigenetic and transcriptional changes. **(A)** FVB/n mice implanted orthotopically with 10^3^ 2.1.1^syn_Luc^ cells and treated with either vehicle (20% HpBCD) or SCH772984 (35 mg/kg), with luminescence reported as mean and standard deviation, N=10 mice per group. **(B, C)** Representative micrographs **(B)** and immunoblots **(C)** of orthotopic tumors treated with vehicle (20% HpBCD) for 48 hours, SCH772984 (35 mg/kg) for 48 hours, or SCH772984 (35 mg/kg) for four weeks. The immunoblots are performed with two biologic replicates, and the micrographs are representative of two biologic replicates. **(D)** Left, hierarchical clustering of RPPA protein expression changes in MIA PaCa-2 cells treated with SCH772984 (1μM) relative to DMSO (1:1,000), performed in triplicate, with individual proteins annotated by cellular function and clusters indicated by circled numbers. Right, restrictive cubic splines of relative growth rate (total doublings per day) of SCH772984-treated cells compared to DMSO-treated cells (top), as well as selected protein expression changes within MAPK and PI3K/AKT/mTOR signaling pathways (middle) and cell cycle/translation markers (bottom). **(E)** Venn diagram depicting total H3K27ac peaks gained or lost (FDR <0.1) in MIA PaCa-2 cells treated with SCH772984 (1 μM) compared to DMSO (1:1,000) for either one week (early) or eight weeks (stable resistance), performed in duplicate. **(F)** Scatter plot comparing gene expression changes in MIA PaCa-2 cells treated with SCH772984 (1 μM) compared to DMSO (1:1,000) for either one week (early) or eight weeks (stable resistance), performed in triplicate. Each dot represents a single gene, with colored dots representing statistically significant (p <10^-3^) gene expression changes at the indicated time points, with statistical significance determined by Wald test using the Benjamini and Hochberg method to correct for multiple hypothesis testing.

To more comprehensively delineate the impact of ERK/MAPK blockade on cellular signaling networks, we utilized reverse phase protein array (RPPA) (*34*) to serially profile *KRAS*-mutant pancreatic cancer cells exposed to dual-mechanism ERK inhibition over a time course ranging from one hour to eight weeks (Figure 1D, Table S1). This array probed diverse RTKs and their associated survival pathways, as well as markers of translational control, pro- and anti-apoptotic regulation, cell cycle control, cytoskeletal dynamics, autophagy, transcription factor activation, and histone modifications. Unsupervised clustering revealed three distinct patterns of protein expression that characterized acquired resistance. Cluster 1 was upregulated during an initial period of stunted growth, but then downregulated as stable resistance developed, and was highly enriched for nodes within the MAPK and PI3K/AKT/mTOR signaling pathways. In contrast, Cluster 2 demonstrated reciprocal downregulation during the initial period of drug sensitivity, followed by a delayed return to baseline expression as resistance developed; this cluster was enriched for markers of cell cycle entry and cap-dependent mRNA translation. These translational regulators represent an established convergence point of integrated ERK/MAPK and PI3K/AKT/mTOR signaling (*35*), and the asymmetric expression patterns between these clusters suggest that intrinsic feedback events are rendered ineffective in the setting of sustained pathway inhibition. The largest cluster (Cluster 3) contained most RTKs and their alternative downstream signaling proteins, which were generally unperturbed by initial drug exposure, and only modestly altered as resistance emerged. Taken together, these findings suggest that pathway reactivation is an endogenous response to ERK/MAPK inhibition that may support early survival, but is insufficient to confer stable resistance in the setting of treatment with a dual-mechanism ERK inhibitor. Furthermore, among diverse alternative signaling pathways probed, none obviously replaced MAPK signaling to drive growth, suggesting a distinct mechanism of resistance to sustained inhibition.

Recent reports have postulated that exposure to kinase inhibitors induces complex changes to the transcriptional and enhancer landscape permitting a drug-tolerant state, and that early changes (within the first week) reflect the necessary adaptations for stable resistance (*21, 22*). However, we have consistently observed stunted cell growth beyond this initial period, and have found that eventual outgrowth is a gradual rather than immediate process, during which cell populations display increasing fitness as stable resistance evolves. This suggests that additional transcriptional evolution may be required to permit the terminally resistant state. In fact, our RPPA analysis demonstrated that all histone markers underwent dynamic changes throughout the adaptive resistance process (Figure S1F), further supporting longer-term transcriptional reprogramming as a potentially necessary component of the terminally resistant phenotype.

To test this hypothesis, we performed RNA sequencing in parallel with chromatin immunoprecipitation-DNA sequencing (ChIP-seq) for acetylated histone 3, lysine 27 (H3K27ac) in treatment-naïve *KRAS*-mutant pancreatic cancer cells, as well as at one week of dual-mechanism ERK inhibition and following the development of stable resistance. H3K27ac represents a histone modification that is associated with transcriptional activation and marks active enhancers; thus, changes in H3K27ac density may broadly signify epigenomic remodeling as cells adapt to environmental stress (*36*). To that end, we found surprisingly limited overlap of H3K27ac peaks gained or lost in the early response compared to stable resistance, counter to previously proposed models suggesting that enhancer remodeling plateaus and is sufficient for resistance by 72 hours of drug exposure (*21, 22*) (Figure 1E, Table S2). And while RNA sequencing revealed a broad early transcriptional response— including well established transcription factors (e.g. EGR1, JUN, KLF2 and FOS), MAPK regulators (e.g. SPRY1/2/4, SPRED1/2), and pro-survival genes in NF-κB/interferon, TGF-β, and alternative tyrosine kinase families (*21, 22*) (Figure S1G)—extensive further transcriptional changes were present in stably resistant cells, and the intersection of dysregulated transcripts between these time points was quite limited (Figure 1F, Table S3). This pattern was consistent among even the most highly up- or downregulated transcripts, including a subset of gene expression changes with opposing directionality at the two time points (Figure S1H). Gene set enrichment analysis (GSEA) further confirmed that there were very few enriched pathways shared between the early response and stable resistance (Figure S1I, Table S4). Taken together, these findings demonstrate an adaptive transcriptional process that is buttressed by parallel remodeling of active enhancers, and that counter to prior models, achieving stable, population-level resistance may require chromatin and gene expression changes that continue to evolve beyond the intrinsic early response to drug exposure. This model is also consistent with recent work proposing that even treatment-naïve cells upregulating key resistance markers require longer-term transcriptional adaptations to cultivate stable resistance (*37*), and prompted us to explore whether targeting key events in transcriptional control might represent an effective strategy for preventing acquired resistance to sustained ERK/MAPK inhibition.

Transcriptionally-mediated resistance programs are broadly vulnerable to manipulation of the epigenetic machinery

To first confirm the generalizability of this long-term transcriptional response to ERK/MAPK suppression, we developed and profiled two additional models of evolved resistance to dual-mechanism ERK inhibition in *KRAS*-mutant cancer cells of distinct tissue origin (colon and lung adenocarcinoma). As in pancreatic cancer cells, these cells also underwent broad early transcriptional changes followed by extensive further adaptations during the development of stable resistance (Figure 2A, Table S3). Notably, the transcriptional programs giving rise to stable resistance demonstrated limited overlap between cells of different tissue origin (Figure 2B). Correspondingly, a complete absence of enriched or suppressed GSEA pathways were shared between resistant models (Figure 2C, Table S4). To test whether these differences were not due simply to tissue type, we developed stable resistance in an additional *KRAS*-mutant pancreatic cancer cell line; indeed, the intersection of differentially expressed genes was no greater among the pancreatic cancer lines than between the other *KRAS*-mutant cancer types (Figure S2A, Table S3), nor were the associated gene annotations using GSEA (Figure S2B, Table S4). Collectively, these findings demonstrate that diverse *KRAS*-mutant cancer cell models undergo large-scale transcriptional changes during sustained ERK/MAPK inhibition, yet the transcriptional programs associated with terminal resistance are heterogenous and model-specific.

**Fig 2.**
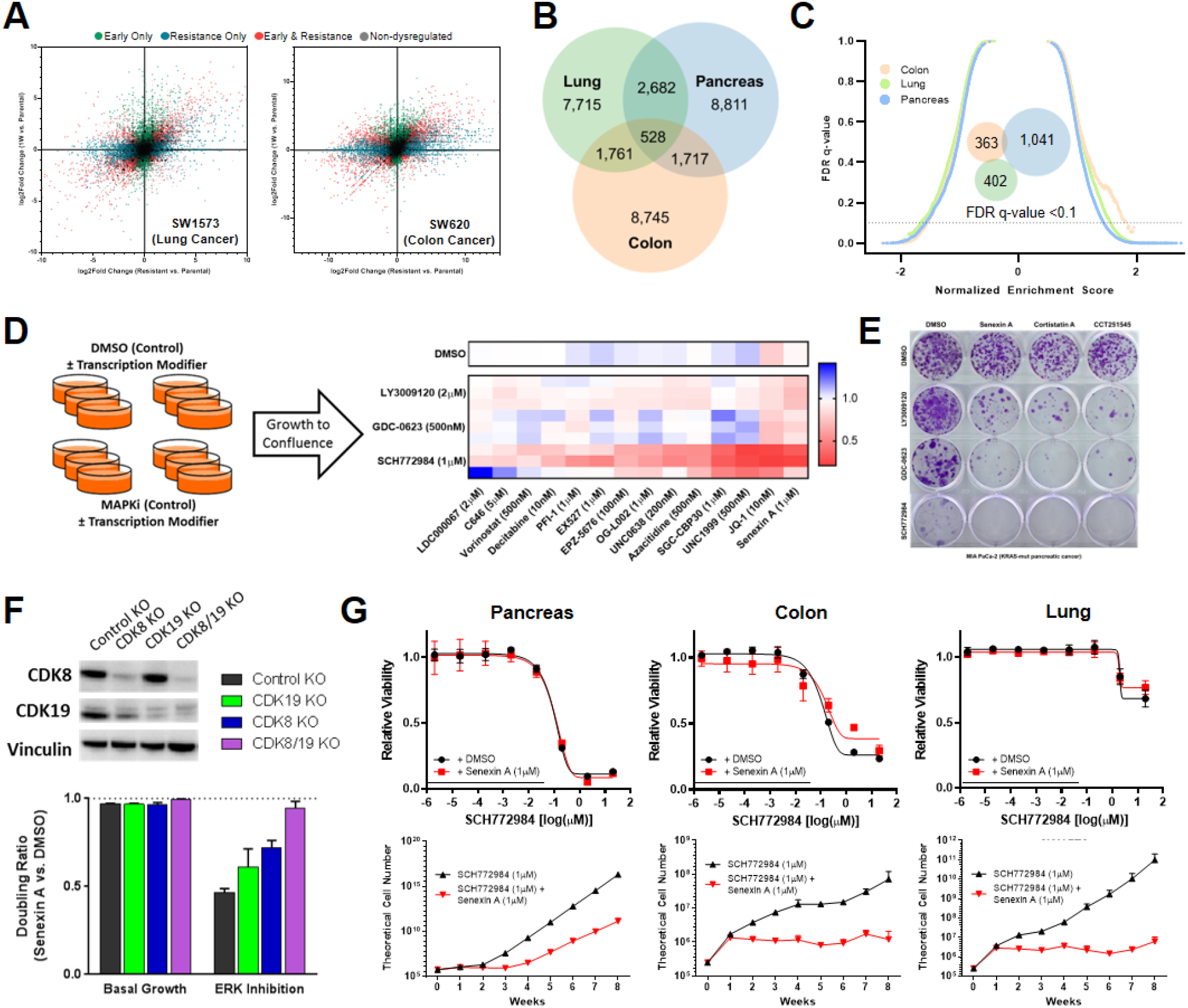
Inhibition of CDK8/19 and other epigenetic modifiers prevents resistance to ERK/MAPK inhibition. **(A)** Scatter plots comparing gene expression changes in SW1573 cells (left) and SW620 cells (right) treated with SCH772984 (1μM) compared to DMSO (1:1,000) for either one week (early) or eight weeks (stable resistance), each performed in triplicate. Each dot represents a single gene, with colored dots representing statistically significant (p <10^-3^) gene expression changes at the indicated time points, with statistical significance determined by Wald test using the Benjamini and Hochberg method to correct for multiple hypothesis testing. **(B)** Venn diagram depicting differentially expressed genes (p <10^-3^) in MIA PaCa-2, SW1573, and SW620 cells treated with SCH772984 (1μM) relative to DMSO (1:1,000) for eight weeks (stable resistance). C) Reverse volcano plot depicting gene set enrichment analysis gene sets from the MsigDB Biologic Process Ontology based on the gene expression data from Figure 2B; Venn diagram depicting enriched gene sets with an FDR <0.1 (insert). **(D)** Schematic representation of the transcriptional modifier pharmacologic screen (left). The heatmap (right) demonstrates the ratio of cell doublings of MIA PaCa-2 cells co-treated with the indicated epigenetic modifier and MAPK inhibitor compared to that MAPK inhibitor alone for three weeks, with each row representing a single biologic replicate. **(E)** Crystal violet staining of 21-day colony growth in MIA PaCa-2 cells treated with the indicated drug combinations, performed in triplicate. **(F)** Immunoblot of MIA PaCa-2 cells with indicated genetic modifications (top); the doubling ratio of those same populations treated with Senexin A (1 μM) relative to DMSO (1:1,000) for two weeks, either alone (basal growth) or in the presence of SCH772984 (1 μM) (bottom), each condition performed in triplicate. **(G)** Eight-point growth inhibition assay of MIA PaCa-2 cells (left), SW620 cells (middle), and SW1573 cells (right) treated with increasing concentrations of SCH772984 in the background of Senexin A (1 μM) or DMSO (1:1000) for four days (top), each condition performed in triplicate; TTP assay of those same cell lines and drug conditions for eight weeks of treatment, each condition performed in triplicate.

Given the obvious challenge of developing strategies to comprehensively target these diverse terminal resistance programs, we instead sought to test whether combining ERK/MAPK inhibition with drugs targeting the epigenetic and transcriptional machinery might broadly perturb the heterogeneous responses to drug exposure. To accomplish this, we performed long-term pharmacologic screens assessing the effects of each member of a panel of drugs targeting diverse transcriptional processes on the development of resistance to second-generation inhibitors of RAF, MEK, and ERK. Inhibitors were selected at doses previously described to achieve target inhibition without significant growth suppression.

In total, we screened 42 combination therapies (Figure 2D, Table S5), revealing, most strikingly, that resistance to sustained ERK inhibition was broadly susceptible to manipulation of the transcriptional machinery, including drugs co-targeting CREBBP/EP300 (SGC-CBP30), EZH1/EZH2 (UNC1999), BET family bromodomain proteins (JQ-1), and CDK8/19 (Senexin A). In keeping with our previous findings of pathway reactivation driving resistance to RAF and MEK inhibition, these targets were generally less vulnerable to combination therapy with transcriptional manipulation. We found that only the BET bromodomain inhibitor JQ-1 and the CDK8/19 inhibitor Senexin A delayed resistance to inhibition at all three ERK/MAPK nodes, with the strongest effect in combination with ERK inhibition. The effect of BET bromodomain inhibition on drug response has been broadly reported (*38–41*), including the long-term interaction of the MEK inhibitor trametinib and the BET bromodomain inhibitors JQ-1 and I-BET151 (*22*), and more recently, the heightened sensitivity to BET inhibition in cells lacking CDK8 and CDK19 (*42*). CDK8 and CDK19 are paralogous proteins that reversibly associate with the multiprotein Mediator complex (*43*), and have never, to our knowledge, previously been implicated in resistance to ERK/MAPK inhibition. CDK8/19 inhibition alone had no effect on basal growth conditions (Figure S2C), nor did it demonstrate short-term synergy with ERK/MAPK inhibition (Figure S2D), suggesting that the impedance of resistance occurred through later-stage inhibition of the adaptive process. Given the profound effect of Mediator kinase inhibition on the activity of drugs targeting all three ERK/MAPK nodes, we chose to characterize this interaction across the three *KRAS*-mutant tissues types for which we had developed models of transcriptionally-mediated resistance.

Over the past decade, an increasing role for Mediator kinase in human cancers has been described, including its function as a colorectal cancer oncogene (*44*), a regulator of super enhancer associated genes in acute myeloid leukemia (*45*), and an effector of melanoma progression (*46*). This has prompted the clinical development of strategies targeting CDK8/19, and we first validated the interaction of combined ERK/MAPK and Mediator kinase inhibition using three structurally distinct preclinical compounds (Senexin A, Cortistatin A, and CCT251545) in long-term colony forming assays at doses shown to selectively inhibit CDK8/19 (*45, 47, 48*). All three compounds profoundly inhibited clonal outgrowth during sustained treatment with RAF, MEK, and ERK inhibition, and again had minimal effect on basal cell growth (Figure 2E). Most notably, combined ERK and CDK8/19 inhibition completely prevented the emergence of resistant colonies at four weeks, supporting a dominant role for Mediator kinases in transcriptional reprogramming during sustained ERK inhibition.

As our work to this point exclusively relied on pharmacologic inhibition of CDK8/19, we next sought to further probe the mechanistic role of each Mediator kinase. CDK8 and CDK19 each possess enzymatic activity, but the proteins can also serve scaffold functions (*49*), and thus protein depletion and kinase inhibition have distinct cellular effects (*50, 51*). To further test whether pharmacologic inhibition was in fact specific to Mediator kinases, we utilized dual CRISPR/Cas9 constructs (*52*) to create knockout derivatives of both CDK8 and CDK19 (sgCDK8/sgCDK19), CDK8 only (sgCDK8/sgControl), CDK19 only (sgCDK19/sgControl), or double-sham control knockouts (sgControl/sgControl) in three different *KRAS*-mutant cancer cell lines. Each condition was then subjected to treatment with DMSO, ERK inhibition, CDK8/19 inhibition, or combined ERK and CDK8/19 inhibition (Figure 2F, Figure S2E,F). In basal growth conditions, treatment with CDK8/19 inhibition had a limited effect on growth in all four derivatives, which was expected given its limited effect in parental cells. In the presence of ERK inhibition, however, CDK8/19 inhibition profoundly suppressed outgrowth of control cells expressing both CDK8 and CDK19, and modestly blocked growth in cells with individual knockout of CDK8 or CDK19, yet had no effect in cells depleted of both CDK8 and CDK19 (Figure 2F). This confirmed that CDK8 and CDK19 were indeed the targets of kinase inhibition responsible for the long-term impedance of resistance.

Finally, to evaluate the effect of CDK8/19 inhibition on sustained MAPK suppression within a more clinically meaningful time course, we utilized an established time-to-progression model (*20, 41, 53*) to test growth in the presence of drug(s) for up to eight weeks. Despite the diverse transcriptional resistance programs observed across tissue types, CDK8/19 inhibition completely prevented lung and colon cancer cells from developing resistance to ERK inhibition for up to eight weeks (Figure 2G, bottom), and markedly delayed the emergence of resistance in pancreatic cancer cells. This consistent finding suggested that CDK8/19 inhibition prevents the establishment of diverse transcriptional programs that drive stable resistance. Of note, combined ERK and CDK8/19 inhibition demonstrated no short-term synergy in any of the cell lines (Figure 2G, top), and CDK8/19 inhibition alone had a negligible effect on long-term basal growth (Figure S2G), positioning this combination as a promising strategy for preventing long-term acquired resistance in *KRAS*-mutant cancers.

### Antagonization of a conserved response network paralyzes further adaptive potential

Given that *KRAS*-mutant cancer cells of distinct tissue origin demonstrated distinct stable resistance programs to ERK inhibition, yet that these programs were nonetheless universally susceptible to CDK8/19 co-inhibition, we next asked whether there might be a common early response to ERK inhibition vulnerable to this combination treatment. In fact, we found that the expression changes induced by ERK inhibition at one week were highly similar between cell lines of distinct tissue origin (Figure 3A), in stark contrast to the limited intersection of expression changes seen in stable resistance (Figure 2B). This indicated that ERK inhibition induces a conserved early transcriptional response before cells undergo further heterogenous adaptations that ultimately establish stable resistance programs. This was further revealed by GSEA (Figure 3B), which demonstrated substantial overlap of enriched gene annotations between tissue types within the early response, again contrasting with the lack of shared pathways in stable resistance (Figure 2C). Broadly, GSEA revealed that this initial response resulted in downregulation of many major anabolic cellular processes such as DNA replication, cell cycle entry, and protein translation (Table S4), which may be necessary for cells to forego growth and replication as unique transcriptional programs are enacted.

**Fig. 3.**
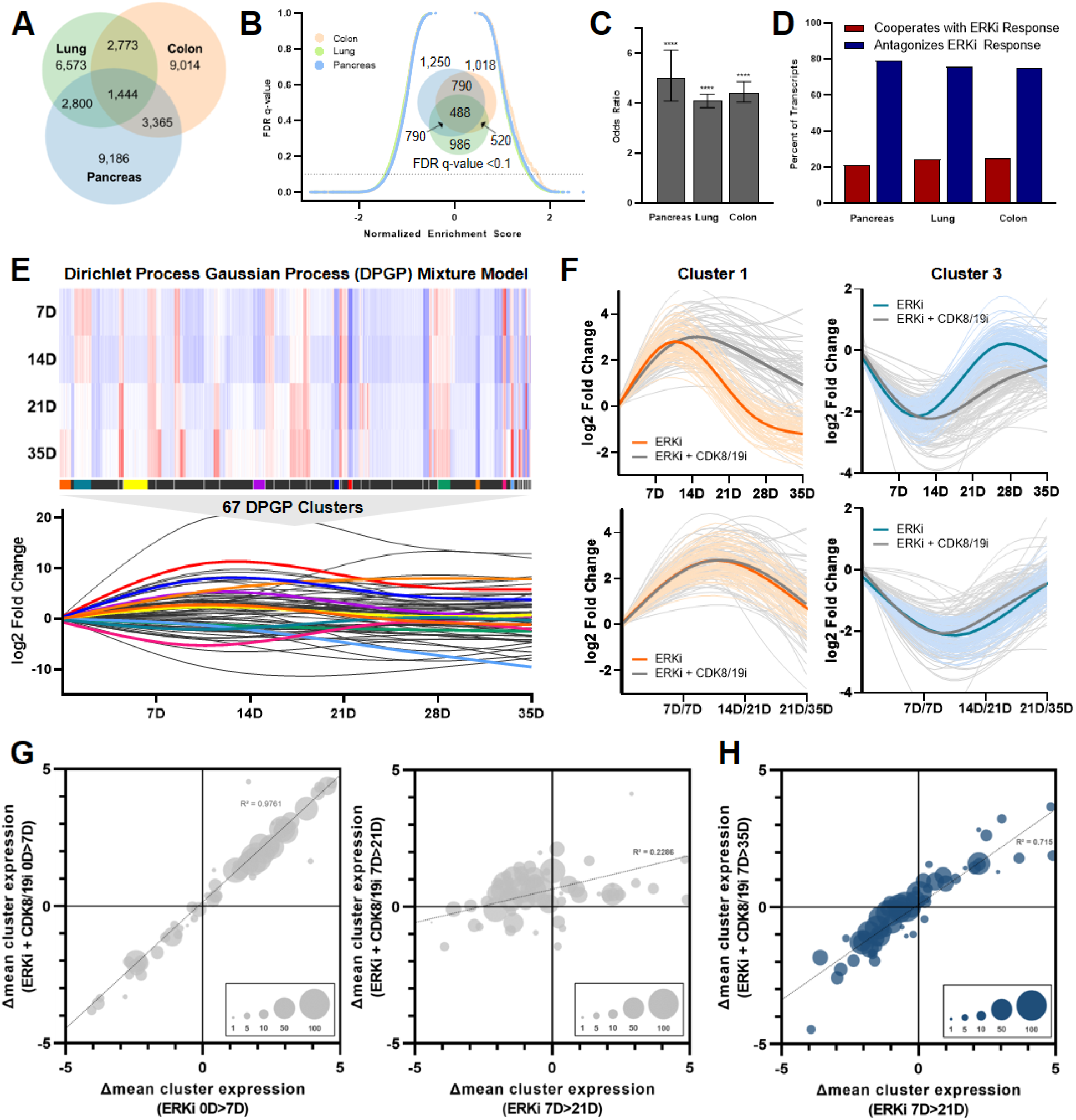
Co-inhibition of CDK8/19 paralyzes long-term transcriptional adaptations by antagonizing the early response to ERK inhibition. **(A)** Venn diagram depicting differentially expressed genes (p <10^-3^) in MIA PaCa-2, SW1573, and SW620 cells treated with SCH772984 (1μM) relative to DMSO (1:1,000) for one week, performed in triplicate, with statistical significance determined by Wald test using the Benjamini and Hochberg method to correct for multiple hypothesis testing. **(B)** Reverse volcano plot depicting gene set enrichment analysis using gene sets from the MsigDB Biologic Process Ontology based on the gene expression data from Figure 3A; Venn diagram depicting enriched gene sets with an FDR <0.1 (insert). **(C)** In MIA PaCa-2, SW1573, and SW620 cells, odds that genes significantly dysregulated by treatment with SCH772984 for one week (1 μM, p <0.001 by Wald statistic) are further up- or downregulated by co-treatment with Senexin A (1 μM), relative to genes that are not significantly dysregulated by treatment with ERK inhibition; p-value calculated according to Sheskin method. **(D)** In these same cell lines, percent of transcripts significantly dysregulated by SCH772984 (1 μM) for which treatment with Senexin A (1 μM) antagonizes or cooperates with these gene expression changes. E) Heatmap (top) depicts differential expression of all genes within the assigned 67 DPGP clusters for MIA PaCa-2 cells treated with SCH772984 (1 μM) relative to DMSO (1:1,000) at the indicated time points; restricted cubic splines of the mean differential expression of each cluster at each time point (bottom). **(F)** Restricted cubic spline of all genes in representative clusters 1 and 3 treated with SCH772984 (1 μM) alone or in combination with Senexin A (1 μM), with the bolded curves representing mean expression changes of each cluster according to treatment condition; the curves on the top reflect equivalent time points for the two treatment conditions, while on the bottom the SCH772984 alone curve is right-shifted by the indicated time intervals. **(G)** Bubble charts reflecting the mean expression changes of each cluster according to treatment condition at one week (left) versus three weeks (right), with the size of each bubble reflecting the number of genes in each cluster. **(H)** Bubble charts reflecting the mean expression changes of each cluster for MIA PaCa-2 cells treated with SCH772984 (1 μM) for three weeks compared to SCH772984 (1 μM) and Senexin A (1 μM) for five weeks, with the size of each bubble reflecting the number of genes in each cluster. ****p<0.0001.

We next asked whether this initial conserved response to ERK inhibition was particularly vulnerable to combined CDK8/19 inhibition. Specifically, we tested whether CDK8/19 co-inhibition preferentially dysregulated the genes identified within this early conserved response network, or whether CDK8/19 inhibition agnostically caused gene expression changes throughout the transcriptome. We found that genes dysregulated by ERK inhibition were significantly more likely to be further altered by combined CDK8/19 inhibition (Figure 3C). We next asked whether this interaction occurred with specific directionality. A nonspecific effect should cause limited expression changes with near-random directionality; alternatively, cooperation or antagonism of a transcriptional program should preferentially amplify or dampen expression changes, respectively. Strikingly, we found that CDK8/19 co-inhibition led to highly organized effects, broadly antagonizing the expression changes within the early conserved response to ERK inhibition (Figure 3D). Notably, despite a highly specific and organized effect on the conserved response to ERK inhibition, the magnitude of antagonization tended to be modest (Figure S3A). Moreover, at this early time point, cells treated with ERK inhibition alone or in combination with CDK8/19 inhibition were phenotypically similar, with equivalent growth inhibition in both treatment conditions (Figure S3B). This led us to ask whether the ultimate consequence of Mediator kinase co-inhibition was the downstream impairment of further transcriptional adaptations necessary for stable resistance.

To do this, cells treated with ERK inhibition alone or combined with CDK8/19 inhibition were serially profiled by RNA sequencing over the five-week time course during which resistance developed (Table S6). Notably, we found that individual transcripts underwent dynamic alterations following the early conserved response, refining their population-level expression throughout this process, including thousands of genes that were both significantly upregulated and downregulated at different time points throughout this evolved process (Figure S3C). Notably, transcriptome-wide differences between treatment conditions became increasingly pronounced over time (Figure S3D), further suggesting that the phenotype induced by CDK8/19 co-inhibition was caused by disruption of this downstream evolutionary process.

In order to visualize, quantify, and evaluate patterns of whole-transcriptome evolution over time, we utilized an established Dirichlet process Gaussian process (DPGP) mixture model, which facilitates time series cluster measurement of genomic features such as gene expression (*54*). Applying this model to the genes with the greatest variance over the course of acquired resistance to ERK inhibition, we identified 67 gene sets reflecting diverse expression trajectories (Figure 3E; Table S7). Like the gene-level analysis between treatment conditions, cluster trajectories between treatment conditions demonstrated overall divergence after the early conserved response (Figure S3E). By quantifying and comparing all cluster-level interval changes, we found that Mediator kinase inhibition exerted its dominant effect following this early response, paralyzing transcriptional reprogramming between weeks one and three of treatment, at which point dually-treated cells resumed a trajectory that mirrored cells treated with ERK inhibition alone (Figures 3F-H), findings we confirmed at the gene level (Figure S3F). By visualizing individual clusters and then shifting the timescale of dually-treated cells to “remove” this period of transcriptional stagnation, the trajectory curves superimposed upon one another (Figure 3F), and we ensured that all cluster trajectories could be realigned with this time shift (Figures 3G,H). We further extrapolated these findings to the entire transcriptome at both the gene-level and using GSEA (Figures S3G,H). Taken together, these findings indicate that the emergence of resistance coincides with transcriptional escape from the early conserved response, a concept that can be observed at the transcript level, using functional annotated gene sets, and using a DPGP mixture model. Co-targeting Mediator kinase therefore antagonizes the conserved response to ERK inhibition, and then exerts its phenotypic effect by preventing further transcriptional changes that are necessary to establish stable resistance.

### Transcriptionally-mediated acquired resistance is driven by distinct terminal mechanisms in different models

Given the magnitude and heterogeneity of gene expression changes observed in each resistant model (Figure 3A,B), yet the ability of combined CDK8/19 inhibition to broadly curtail the emergence of resistance across models (Figure 2G), we next sought to map out a specific resistance mechanism in one of these models. To date, efforts to characterize the function of individual components of a transcriptional program have been limited by the scalability of candidate-based approaches. We considered it unrealistic to implicate the functional importance of individual genes based simply on expression changes, as some highly dysregulated transcripts were likely secondarily regulated passengers of alterations to the chromatin architecture, vestigially over- or underexpressed following the early response, or simply dysregulated at random.

In order to test the consequence of gene-level transcriptional events, we designed a specialized loss-of-function library of CRISPR/Cas9 constructs targeting the subset of genes most dysregulated throughout the adaptive process in MIA PaCa-2 cells (Figure 4A, Table S8). Criteria for library selection included the 200 genes most up- or downregulated at early drug exposure and at stable resistance, with priority based on the differential expression level and significance across two biological replicate conditions, each performed in technical triplicate (all 716 selected genes meeting these criteria had an absolute log2 fold change of >0.5 and a p-value <1.0 x 10^-5^ in both models; Table S9). Also included were 100 control genes selected for their general essentiality or dispensability (*55*), 50 internal control genes which exhibited minimal expression changes during the adaptive process, and 50 non-targeting control guides. The library was cloned into an established lentiviral system (*56*), and all sgRNA sequences (five guides per genes and non-targeting controls) were selected from a reputable genome-wide library (*57*). Drug-naïve and resistant cells were then transduced with lentivirus, subjected to puromycin selection, and sampled serially in both drug treatment and vehicle control conditions in order to define genes important for drug sensitivity in both the naïve and resistant states. The composition of the sgRNA pools was determined by deep sequencing, taking the average of all five constructs to produce a gene-level score across biologic replicates for each condition. We validated our approach by comparing final and initial sgRNA pools from treatment-naïve parental cells across replicates, focusing on known essential genes, non-essential genes, and non-targeting controls (Figure 4B).

**Fig. 4.**
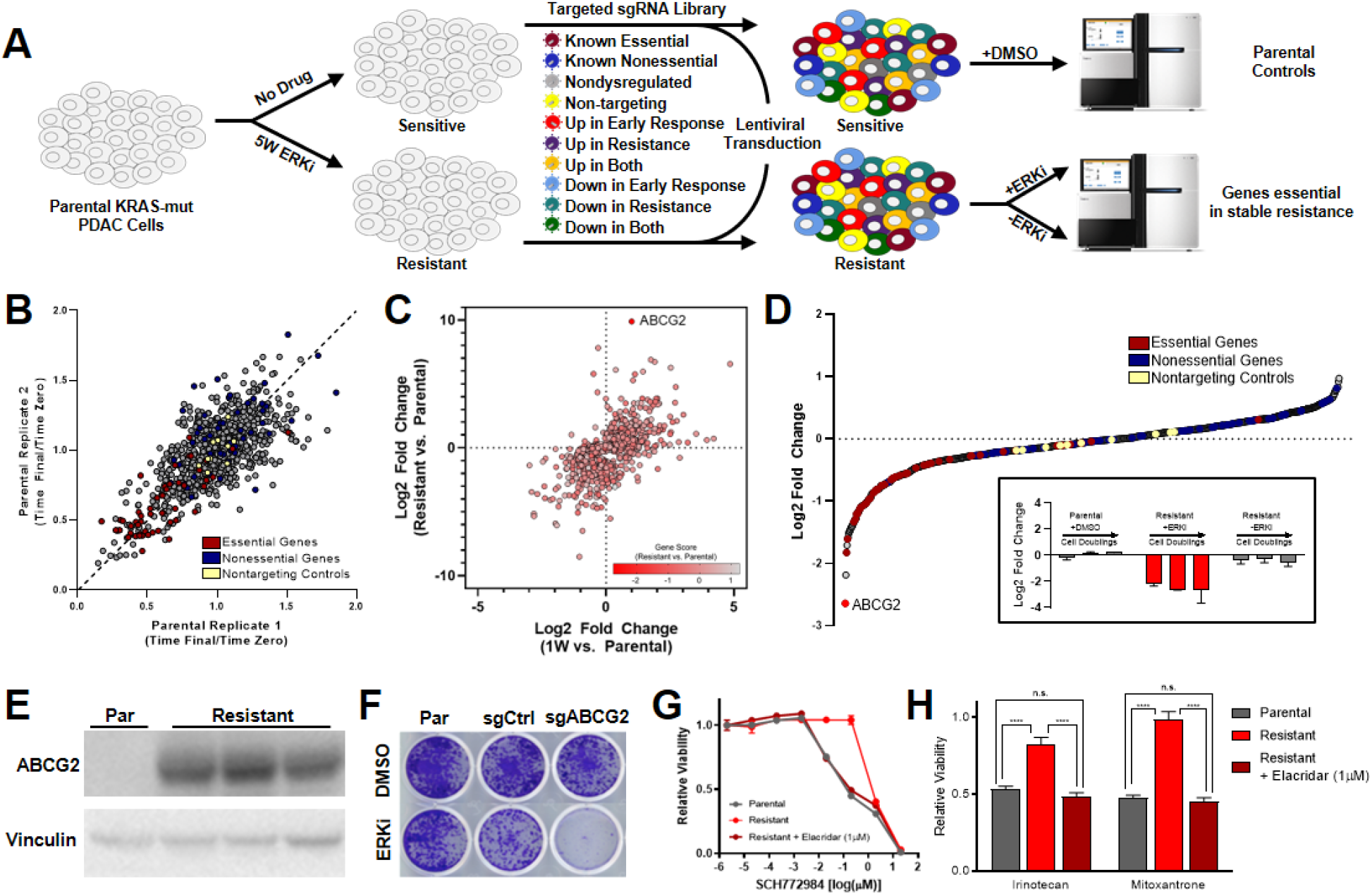
Transcriptional plasticity permits diverse mechanisms of resistance to ERK/MAPK inhibition. **(A)** Schematic depicting CRISPR/Cas9 loss-of-function screen. **(B)** Replicate-to-replicate comparison of gene-level essentiality phenotypes in MIA PaCa-2 cells treated with DMSO (1:1,000) alone. Essential controls are shown in red, non-essential controls in blue, and non-targeting controls in yellow. **(C)** Scatter plot of genes included in loss-of-function screen comparing gene expression changes in MIA PaCa-2 cells treated with SCH772984 (1 μM) compared to DMSO (1:1,000) for either one week (early) or eight weeks (stable resistance), with loss-of-function gene score indicated by color gradient. Gene score is calculated as the log2 fold change of the fractional representations (time final/time zero) of resistant cells treated with SCH772984 (1 μM) compared to parental cells treated with DMSO (1:1,000) at the screen midpoint (19 days). **(D)** Gene-level representation of essential phenotypes in evolved resistant MIA PaCa-2 cells treated with SCH772984 (1 μM) at the screen midpoint (19 days), ranked by their mean log2-transformed gene score across duplicates; the insert shows the effect of ABCG2 loss in parental cells treated with DMSO (1:1,000, left), resistant cells treated with SCH772984 (1 μM, middle), and resistant cells treated with DMSO (1:1000, right) at all three screen time points (15, 19, and 23 days). **(E)** Immunoblot of treatment-naïve parental MIA PaCa-2 cells and three independently evolved resistant derivatives of this same cell line to SCH772984 (1μM). **(F)** Crystal violet staining of MIA Paca-2 cells evolved resistant to SCH772984 (1μM) with either no alteration, control knockout, or ABCG2 knockout, treated with either SCH772984 (1 μM) or DMSO (1:1,000) for one week; representative photograph of all conditions performed in triplicate. **(G)** Eight-point growth inhibition assay of parental and evolved resistant MIA PaCA-2 cells treated with increasing concentrations of SCH772984, performed in triplicate; one triplicate set of evolved resistant cells is treated with the ABCG2 inhibitor Elacridar (1 μM) in the background. **(H)** GI50 values derived from eight-point dose-response curves of parental and evolved resistant MIA PaCA-2 cells treated with increasing concentrations of the indicated chemotherapies, with and without Elacridar (1 μM).

These results revealed that while extensive transcriptional changes were necessary to achieve a stably resistant phenotype, the vast majority of these genes were not independently necessary for the maintenance of the resistant state, as their knockout failed to re-sensitize resistant cells to ERK inhibition (Figure 4C, Figure S4A). However, the knockout of one gene selectively upregulated in resistant cells—encoding the ATP-binding cassette protein ABCG2, a well-known multidrug transporter with broad polysubstrate specificity and normal physiologic functions in uric acid efflux and xenobiotic export (*58*)—dramatically sensitized resistant cells to ERK inhibition (Figures 4C, D). ABCG2-mediated resistance was extensively validated by demonstrating its reproducible upregulation, and subsequently that both gene knockout and pharmacologic inhibition of its ATPase function fully reversed ERK inhibitor resistance, while sensitizing cells to treatment with other known ABCG2 substrates (Figures 4E-H). ABCG2 loss demonstrated no functional consequence in basal growth or in resistant cells when drug was removed (Figure 4D (insert), Figures S4A, B), confirming it as a *bona fide* driver of drug resistance in this cell model. To ensure that ABCG2-upregulation was not simply occurring via the outgrowth of small population of ABCG2 high-expressing “persister” cells, we demonstrated that the vast majority of individual cells were, in fact, capable of developing resistance across a consistent timescale (Figure S4C). Interestingly, the reproducible ABCG2-mediated resistance observed in MIA PaCa-2 cells (Figure 4E) was not observed in *KRAS*-mutant colon and lung cancer models, which evidenced neither *ABCG2* upregulation (Figure S4D) nor resensitization to ERK inhibitor by pharmacologic ABCG2 inhibition (Figure S4E). Collectively, these findings demonstrate that transcriptional *ABCG2* upregulation reproducibly drives acquired resistance to ERK inhibition in MIA PaCa-2 cells, but that differing mechanisms drive acquired resistance in other cell line models, highlighting the notion that strategies targeting transcriptional adaptation are likely to be more broadly effective at blocking resistance than strategies targeting discrete gene expression changes.

### Co-targeting Mediator kinases delays resistance to ERK inhibition across translational models of *KRAS*-mutant cancers

To assess the translational generalizability of combined ERK and Mediator kinase inhibition, we evaluated this strategy across multiple *in vitro* and *in vivo* models of *KRAS*-mutant cancer, including additional eight-week time-to-progression models, patient-derived rectal cancer tumoroids, and a GEMM-derived orthotopic, syngeneic mouse model of pancreatic cancer.

We first expanded upon the existing time-to-progression model by testing the combination of ERK and CDK8/19 inhibition across a panel of diverse *KRAS*-mutant pancreatic, lung, and colorectal cancer cell lines. In all models, cotreatment with CDK8/19 inhibition delayed the emergence of resistance, in many cases completely preventing resistance until the assay was terminated at eight weeks (Figure 5A, S5A). Again, combined ERK and CDK8/19 inhibition demonstrated no short-term synergy in any of the cell lines (Figure S5B), and CDK8/19 inhibition alone had a negligible effect on long-term basal growth (Figure S5C). In addition, we tested two lines with distinct oncogenic drivers (both *KRAS* wildtype; one *BRAF* mutant melanoma (A375) and one *EGFR* driven colorectal cancer (LIM1215)) with respective genotype-specific targeted inhibitors. In these two models, CDK8/19 inhibition had no effect on the emergence of resistance (Figures 5A, S5A). Notably, resistance to these paired oncogenic drivers/targeted agents typically develops via genetic pathway reactivation (*59, 60*). Thus, the ability of CDK8/19 inhibition to forestall resistance may require transcriptionally-mediated resistance mechanisms, and thus may serve as an effective combination strategy in other contexts where resistance develops via large-scale, long-term transcriptional adaptations.

**Fig. 5.**
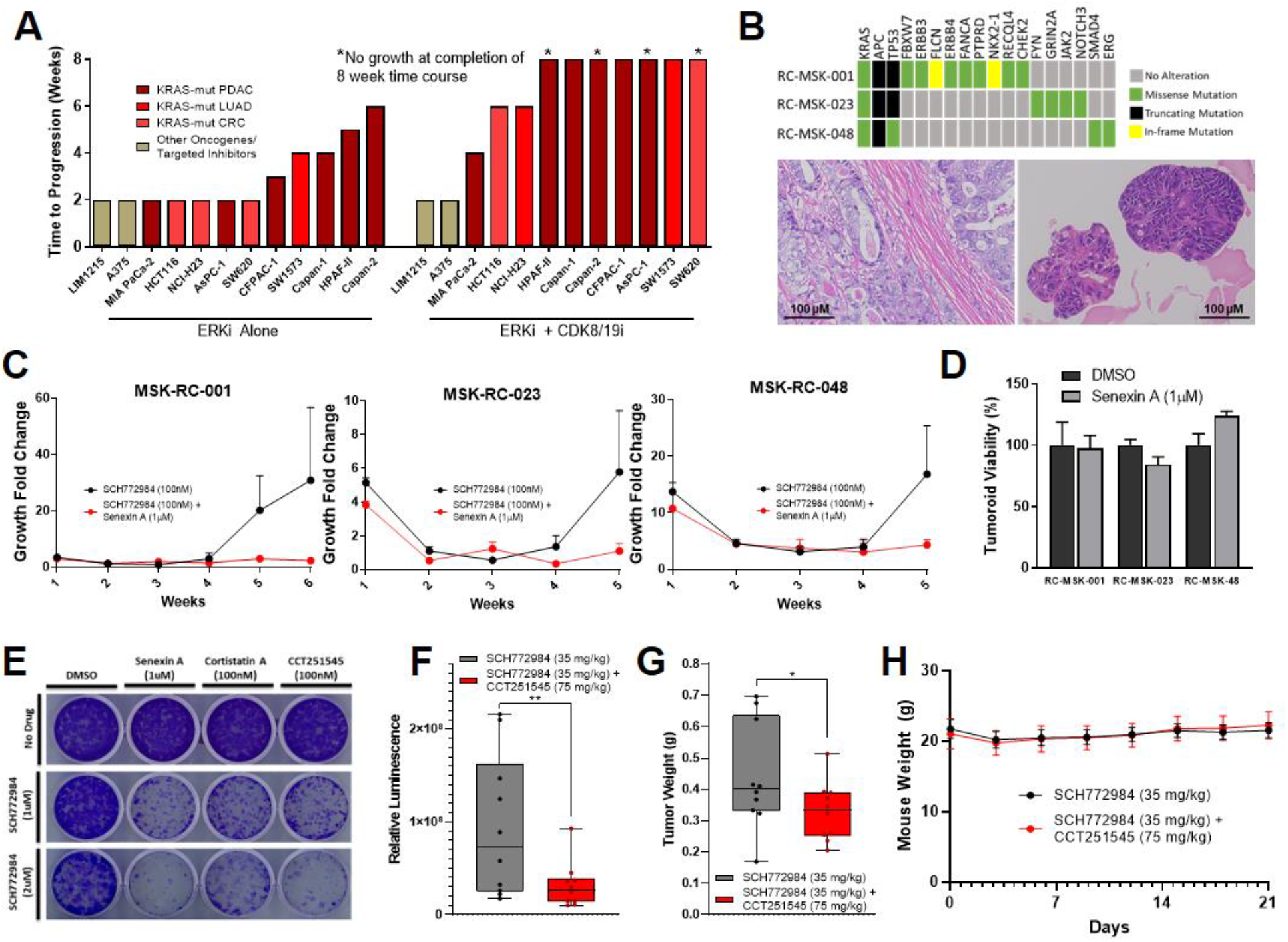
Cotargeting Mediator kinase prevents resistance to ERK/MAPK inhibition. A) Bar chart depicting the time at which resistance emerged to treatment with either SCH772984 (1 μM) alone or SCH772984 (1 μM) in combination with Senexin A (1 μM). **(B)** Oncoplot of three rectal cancer tumoroids based on MSK-IMPACT testing (top), with representative micrographs below of RC-MSK-001 patient tumor (left) and its corresponding tumoroid (right). **(C)** TTP assays for rectal cancer tumoroids treated with SCH772984 (100nM) alone or in combination with Senexin A (1 μM). **(D)** Relative viability of rectal cancer tumoroids treated with Senexin A (1 μM) compared to DMSO (1:1,000) for 14 days. **(E)** Crystal violet staining of 14 day colony growth of 2.1.1^syn_Luc^ cells treated with the indicated drug combinations. **(F,G)** Box plot demonstrating whole body luminescence **(F)** and tumor weights **(G)** after orthotopic implantation of 10^3^ 2.1.1^syn_Luc^ cells into FVB/n mice and treatment with SCH772984 (35 mg/kg) in combination with or without CCT251545 (75mg/kg) for 21 days, 10 mice per group. **(G)** Mean mouse weights throughout treatment described in Figure 5F, G.

Next, we tested the combination of ERK and CDK8/19 inhibition in three *KRAS*-mutant, patient-derived tumoroid models of rectal cancer (Figure 5B-D). These tumoroids had previously been shown to retain the molecular, histological, and clinical features of the patient tumors from which they were derived, as well as accurately predict individual patient responses to chemotherapy and radiation therapy (*61*). Further, all tumoroids had undergone genomic profiling via MSK Integrated Mutation Profiling of Actionable Cancer Targets (MSK-IMPACT) (Figure 5B) (*62*), which demonstrated 92% and 77% concordance, respectively, with oncogenic and other clonal mutations relative to the tumors from which they were derived. Like the cell line time-to-progression models, tumoroids developed resistance to ERK inhibition within five weeks of treatment (Figure 5C). Alternatively, tumoroids treated with combined ERK and CDK8/19 inhibition demonstrated complete growth suppression up to eight weeks (Figure S5D). Again, inhibition of Mediator kinase alone had no effect on tumoroid growth (Figure 5D).

Finally, we tested the combination of ERK and CDK8/19 inhibition in a well-credentialed, GEMM-derived orthotopic, syngeneic mouse model of pancreatic cancer. We first tested three independent CDK8/19 inhibitors *in vitro* using a cell line derived from the same GEMM. Cell growth was unaffected by each inhibitor alone, yet resistance to ERK inhibition was markedly impeded by co-inhibition of Mediator kinase, with similar efficacy between compounds (Figure 5E). Of these three inhibitors, we selected CCT251545, a selective inhibitor of CDK8/19 (*47, 63*) for *in vivo* studies (Cortistatin A is not yet commercially available, and Senexin A has known limited oral bioavailability). Treatment with ERK inhibition alone dramatically reduced initial tumor growth, yet resistance to treatment developed within two weeks of drug exposure (as in Figure 1A). In contrast, inhibition of ERK and CDK8/19 delayed the acquisition of resistance (Figure 5F), resulting in a 34% reduction in tumor weight at study endpoint (Figure 5G) without additional toxicity (Figure 5H). As in the cell line and tumoroid models, inhibition of Mediator kinase alone had no effect on *in vivo* tumor growth or overall toxicity (Figures S5E-G), positioning CDK8/19 co-inhibition as an effective and well-tolerated strategy for preventing resistance to sustained MAPK inhibition.

## Discussion

The RAS family of oncoproteins represent the most frequently mutated genes in human cancers (*1, 2*), and *KRAS*, in particular, is altered in a subset of the most common and deadly solid tumors, including >90% of pancreatic adenocarcinomas (*28*), ~40% of colorectal adenocarcinomas (*29*), and ~30% of lung adenocarcinomas (*30*). While therapies targeting the ERK/MAPK pathway represent an attractive strategy for these malignancies, to date RAS pathway inhibitors have been limited by complex negative feedback loops reactivating endogenous signaling networks (*4, 7*). Newer agents designed to prevent ERK/MAPK pathway reactivation have also proven susceptible to evolved resistance (*17–20*) Given these obstacles, the majority of *KRAS*-mutant pancreatic, lung, and colorectal cancers continue to be treated with conventional cytotoxic chemotherapies (*64–67*).

Here, we demonstrate that sustained ERK/MAPK inhibition causes enhancer remodeling and transcriptional reprograming that is broadly vulnerable to pharmacologic perturbation of chromatin binding and modifying factors (e.g. CBP/p300, BRD4), as well as Mediator kinases. Co-targeting CDK8/19 antagonizes a conserved early response to ERK inhibition, which then paralyzes the additional transcriptional adaptations observed at the population level that appear necessary to establish stable resistance. Across translational models, inhibition of Mediator kinase alone demonstrated only minimal effect on basal cell growth, and long-term inhibition of CDK8/19 was well-tolerated while preventing acquired resistance to ERK inhibition. Thus, this work validates a strategy for robust and potentially nontoxic prevention of ERK/MAPK inhibitor resistance, a problem that to this point has proven recalcitrant.

This study builds on prior work demonstrating that repression of early inhibitor-stimulated transcriptional events can broadly improve therapeutic efficacy without *a priori* knowledge of all potential terminal resistance mechanisms (*21, 22*). This strategy differs from “rational” combination approaches that pair oncogene targeted therapies with drugs inhibiting known resistance mechanisms, which have provided only transient effectiveness in both preclinical studies and patient trials (*8–10*). Notably, our own work using CRISPR/Cas9-based screening to identify co-targets for inhibition with second-generation ERK/MAPK inhibitors in *KRAS*-mutant tumors yielded potent short-term combinations that nonetheless gave rise to resistance; in fact, three-drug combinations leveraging either the primed apoptotic state or the addition of conventional cytotoxic chemotherapies were required to achieve durable responses *in vivo (20*).

Here, we coopt a strategy to broadly impede a conserved gene expression network in order to block further transcriptional adaptations and circumvent an array of potential resistance mechanisms. The distinct transcriptional resistance programs identified in this study highlight the diversity of mechanisms that may be simultaneously targeted by this approach. Interestingly, prior work in this space has suggested that a transient epigenetic state (*27*) or rare cell transcriptional variability (*37*) leads to resistance in only a small subpopulation of cells. In contrast, findings from this study suggest that a large fraction of the tumor cell population may be capable of transcriptional reprogramming in order to support stable resistance programs, although future work is necessary to elucidate the population dynamics and kinetics of these cell-level events.

Notably, in assessing resistance to multiple levels of MAPK inhibition, we found that inhibition of RAF and MEK were substantially less vulnerable to strategies co-targeting the epigenetic machinery than ERK inhibition. This may be secondary to distinct patterns of resistance; signaling analyses suggest that while even newer inhibitors of RAF and MEK seem susceptible to pathway reactivation, second-generation ERK inhibition may sustainably block MAPK signaling at its terminal node. Moreover, among the primary ERK/MAPK pathway proteins, ERK uniquely acts on both cytoplasmic and nuclear targets (*68*). Together, this may result in transcriptional reprogramming as the dominant mechanism of resistance for persistent ERK/MAPK blockade. In fact, ERK has been considered a potential “Achilles’ Heel” in targeted MAPK signaling due to its bottleneck position between MEK and an array of downstream substrates (*69*), and preclinical studies have demonstrated impressive single-agent activity of ERK inhibition across a spectrum of ERK/MAPK-dependent cancer types (*13, 15*). Nonetheless, reports of *in vitro* resistance to second generation, dual-mechanism ERK inhibitors (*19, 20, 70*) may portend disappointing results as these drugs reach clinical trials in human patients (*71, 72*). Thus, strategies to circumvent broad patterns of resistance will be highly useful for further clinical development of ERK inhibitors.

Here, we show that CDK8/19 inhibition universally impairs resistance to ERK/MAPK inhibition in *KRAS*-mutant cancer cells, and that the combination of ERK and CDK8/19 inhibition durably prevents resistance in a panel of translational models. CDK8 and CDK19 are paralogous proteins that reversibly associate with the Mediator complex (*43*), a global regulator of pol II transcription (*73*). Mediator is recruited to enhancers and promoters through interactions with sequence-specific DNA-binding transcription factors (TFs), which represent a major class of proteins phosphorylated by Mediator kinases (*50*). TFs drive all physiological processes, including changes in cell state (*74, 75*), and inhibition of Mediator kinase function has been shown to block stimulus-specific TF activation (*49, 76, 77*). Consequently, inhibition of Mediator kinase function during sustained ERK/MAPK inhibition may impede TF-dependent remodeling of gene expression networks that are required for development of stable resistance to ERK inhibition.

While we found that Mediator kinase co-inhibition robustly suppressed resistance to ERK inhibition in our cell line and organoid models, we observed a shorter period of delayed resistance *in vivo*. It is possible that this reflects the highly aggressive nature of the orthotopic KPC model of pancreatic cancer which we used, and that in less aggressive *KRAS*-mutant models resistance may be delayed in a manner that would be suitable for transitioning to human trials. However, a second hypothesis is that the prevention of resistance *in vivo* was limited by the oral bioavailability and metabolic stability of CCT251545, which has been broadly problematic for CDK8/19 inhibitors to date. This is supported by the strong effect we observed *in vitro* using KPC-derived cells with three separate CDK8/19 inhibitors. Currently, the natural product Cortistatin A is considered among the most promising preclinical compounds in terms of its potency and bioavailability, but it is not yet commercially available, in part due to a complicated synthesis process (*45, 50*). Thus, there is a broad interest in the clinical development of additional potent, bioavailable CDK8/19 inhibitors. As these emerge, future studies will focus on validating our findings across additional *in vivo* models of *KRAS*-mutant cancers.

In sum, this study provides strong support for the therapeutic strategy of combining ERK and Mediator kinase inhibition. Our results, which combined diverse cell lines with tumoroids and *in vivo* models of *KRAS*-mutant cancers, suggest that Mediator kinase inhibitors may be broadly effective in blocking short- and long-term transcriptional changes required for the emergence of drug resistant cell populations. Additional studies testing novel classes of ERK/MAPK pathway inhibitors in combination with drugs that target epigenetic and transcriptional processes may provide an efficient way of overcoming the rapid resistance that has traditionally been observed with genetically-targeted anticancer therapies.

## Materials and Methods

### Cell lines and reagents

All cell lines were grown at 37°C in 5% CO_2_. Human pancreatic, lung, and colorectal cancer cell lines were grown in DMEM/F12, 10% FBS, and 1% penicillin/streptomycin. Mouse cell lines were grown in DMEM, 10% FBS, and 1% penicillin/streptomycin. All cell lines were purchased from American Type Culture Collection (ATCC) or Duke University Cell Culture Facility (CCF). All cell lines were authenticated using Promega PowerPlex 18D kit or were purchased within 6 months from Duke CCF. Drugs were purchased from Selleck Chemicals, except Cortistatin A (a gift from the laboratory of Matthew Shair) and CCT251545 (which was provided by The Institute of Cancer Research).

### Mouse studies

As previously described (*78*), female FVB/n mice were injected orthotopically (into the head of the pancreas) with 1,000 luciferase-expressing p53 2.1.1^syn_Luc^ mouse pancreatic tumor-derived cells (*Kras*^G12D^/*Tp53*^-/-^; provided by Dr. Eric Collisson, UCSF) that had been resuspended in 50% Hanks’ balanced salt solution (Gibco), 50% LDEV-free Matrigel (Corning) at a concentration of 25 cells/mL. Ketamine (80 mg/kg), xylazine (8 mg/kg), and acepromazine (1 mg/kg) were used to anesthetize the mice prior to surgery. A tuberculin syringe with a 30-g needle was inserted into an abdominal incision and used for implantation. The incision was closed using surgical staples. After a one-week recovery, initial IVIS imaging with D-luciferin substrate (Perkin Elmer) was performed using an IVIS Lumina optical imaging system. For each independent experiment, mice were separated into groups of 10, and treated daily with either vehicle (20% HdpCD, IP), SCH772985 (35 mg/kg, IP), CCT251545 (75 mg/kg, OG), or a combination of both SCH772985 (35 mg/kg, IP) and CCT251545 (75 mg/kg, OG). CCT251545 was resuspended in ORA-Plus (Perrigo). IVIS imaging was performed at weekly time points. Mice were sacrificed at the conclusion of the study, the pancreas was removed, and tumors were then excised from the pancreas and weighed.

### Short-term growth inhibition assay

As previously described (*20*), cells were seeded into 96-well plates at 5,000 cells/well. To generate GI50 curves, cells were treated with vehicle using an eight-log serial dilution of drug. For certain experiments, cells were treated in the background of a constant concentration of a second drug or DMSO (1:1,000) to test combinatorial effects. Each treatment condition was represented by at least three replicates. Three days after drug addition, cell viability was measured using Cell Titer Glo (Promega). Relative viability was then calculated by normalizing luminescence values for each treatment condition to control treated wells. Dose-response curves were fit using GraphPad/Prism 6 software.

### Long-term time-to-progression (TTP) assay

As previous described (*41*), cells were seeded at 250,000 per plate in 10 cm plates in triplicate. The next day, drugs were added at indicated concentrations. At weekly time points, cells were counted and replated at 50,000-100,000 per plate, and drug/media was replenished. Virtual cell counts were calculated based upon the number of cells plated, the growth rate, and the final cell counts at each week. For most experiments, the TTP assay was terminated at eight weeks. For the double CRISPR/Cas9 knockout of CDK8 and CDK19, the TTP assay was terminated at two weeks.

### Western blotting and antibodies

Immunoblotting was performed as previously described (*20*). Membranes were probed with primary antibodies (1:1,000 dilution) recognizing KRAS (CST #14429), p-MEK (CST #9121), T-MEK (CST #4694), p-ERK (CST #9101), T-ERK (CST#4695), vinculin (CST#4650), CDK8 (CST #4106), and CDK19 (Sigma HPA007053).

### Immunohistochemistry

Formalin-fixed, paraffin-embedded tissue blocks of mouse orthotopic tumors were sectioned and stained with hematoxylin and eosin, as well as antibodies targeting phospho-MEK (CST #2338), phospho-ERK (CST #4370), and Ki-67 (Sigma SAB5600249). Negative controls were performed on all runs using an equivalent concentration of a subclass-matched immunoglobulin. All immunohistochemistry was performed in biologic duplicate. Images were then digitized and visualized using Aperio Imagescope.

### Reverse-phase protein array sample preparation and analysis

MIA PaCa-2 cells were treated with SCH772084 (1 μM) or DMSO (1:1,000) for the indicated durations at which point plates were washed twice in ice-cold PBS and then frozen at −80 °C. At the completion of all time points, plates were scraped and RPPA analysis was performed as previously described (*34*). All samples were conducted in biological triplicate and normalized to a DMSO control at each respective time point.

### Generation of evolved resistant cell lines

Cells were plated at a density of 250,000 cells in 10 cm plates. The following day, SCH772984 (1 μM) was added. Cells were then split at weekly time points and replated at a density of 50,000, and drug/media was replenished. Lines were considered resistant when the growth rate exceeded 50% of that of its parental, treatment-naïve derivative, and terminal resistance was defined once there was no interval increase in growth rate over a two-week period.

### ChIP-Seq

Cells were grown in 15cm^2^ plates in either SCH772984 (1 μM) or DMSO (1:1,000) in biologic replicate. For the one-week samples, 560,000 and 110,000 cells were plated in the SCH772984 and DMSO conditions, respectively, to achieve 70% confluence at one week. For the terminally resistant cells, evolved resistance was achieved as described above, and then this cell population was plated at 110,000 cells per 15cm^2^ plate in biologic replicate and treated for one additional week with SCH772984. At this point, cells were cross-linked using 1% formaldehyde followed by 2.5M glycine and washed with ice-cold PBS. Cross-linked cells were then scraped in RIPA buffer and snap-frozen. As previously described (*79*), chromatin was sheared and antibody-conjugated (H3K27ac active motif: 39133) Protein A Dynabeads beads (Invitrogen) were used for chromatin extraction. Bound chromatin was eluted, crosslinking was reversed, and DNA was then purified by phenol chloroform extraction. Sequencing libraries were prepared using the KAPA Hyper Prep Kit (KAPA Biosystems). Sequencing was performed on an Illumina NextSeq. For data analysis, peaks were first identified using MACS2 with a configuration suitable to detect narrow peaks as those typically observed in H3K27ac data. A union peakset of all possible acetylation events identified across conditions was then defined. Using this common set, reads in peaks were computed using featureCounts with default parameters. Lastly, to detect differential binding events a negative binomial model using DESeq2 was applied to the counts, followed by a Wald test to compute p-values. Gained peaks represent both regions gaining or increasing acetylation when compared to control samples. Conversely, sites with depleted or decreased signal are referred to as lost peaks.

### RNA sequencing and Gene Set Enrichment Analysis

RNA was extracted from cell pellets using the RNeasy Kit (Qiagen), and library preparation and sequencing was performed by the Duke Sequencing and Genomic Technologies Shared Resource. RNA-seq data was processed using the TrimGalore toolkit which employs Cutadapt to trim low quality bases and Illumina sequencing adapters from the 3’ end of the reads. Only reads that were 20nt or longer after trimming were kept for further analysis. Reads were mapped to the GRCh37v75 version of the human genome and transcriptome using the STAR RNA-seq alignment tool. Reads were kept for subsequent analysis if they mapped to a single genomic location. Gene counts were compiled using the HTSeq tool. Only genes that had at least 10 reads in any given library were used in subsequent analysis. Normalization and differential expression was carried out using the DESeq2 Bioconductor package with the R statistical programming environment. The false discovery rate was calculated to control for multiple hypothesis testing. Gene set enrichment analysis was performed to identify gene ontology terms and pathways associated with altered gene expression for each of the comparisons performed.

### Long-term pharmacologic screens

Cells were plated at 20,000 cells per 10 cm plate in triplicate. The next day, drugs were added at the indicated doses. Media and drug were then exchanged weekly until the control condition for each MAPK inhibitor grew to 90% confluence, at which point cells in all plates in that treatment condition were counted, and population doublings were calculated and compared to the control condition.

### Clonogenic growth assay

Cells were seeded at 2,000-10,000 cells per well. The next day, cells were drugged at the indicated doses. At assay completion, plates were rinsed with PBS and fixed and stained with 0.5% (wt/vol) crystal violet in 6.0% (vol/vol) glutaraldehyde solution (Thermo Fisher Scientifics) for 30 minutes at room temperature. Plates were rinsed in distilled H_2_O and photographed the following day.

### Generation of CRISPR/Cas9 knockout derivatives

CRISPR constructs were cloned into the lentiCRISPR v2 vector as previously described (*80*). After lentivirus production, viral tittering, and transduction (*20*), cells were replated into fresh media in 10 cm plates, and one day later puromycin was added (2μg/mL). Two days later, puromycin was removed, and cells were considered stably transduced five days later. For double knockout experiments, we also used a modified version of the lentiCRISPR v2 vector (lentiCRISPR v2-Hygro) (*52*), in which the puromycin resistance gene was exchanged for a hygromycin resistance gene. This allowed us to sequentially generate dual-knockout derivatives, and transduction was performed as described above, after the first knockout was considered stable.

### Dirichlet Process Gaussian Process (DPGP) Mixture Model

In order to identify gene sets exhibiting similar gene expression trajectories, we applied a previously published DPGP model (*54*). Briefly, DPGP simultaneously models data clusters using a Dirichlet process and dependencies on data timepoints with Gaussian processes. This joint model allows us to identify large and subtle differences in transcriptional profiles over time. Once DPGP clusters were established, changes in gene expression among each component was defined as the log2 fold change relative to DMSO-treated parental cells at each time point. Mean cluster expression was defined as the average log2 fold change of all cluster components at each time point relative to DMSO-treated parental cells.

### Loss-of-function CRISPR/Cas9 screens

Our sgRNA library was designed, cloned and amplified as previously described, as was virus production, titering and transduction (*20*). Treatment-naïve and evolved resistant MIA PaCa-2 cells were separately transduced with library virus at a multiplicity of infection of 0.2 at 1000x coverage, and then these populations underwent seven days of puromycin selection. Following puromycin selection, each population was divided and treated with either SCH772984 (1 μM) or DMSO (1:1,000). Each condition was conducted in biologically independent replicates and carried out at >1,000x coverage for five weeks, and cells were split once they reached 80% confluence. At each split, excess cells beyond those needed to maintain 1000x coverage were pelleted and stored at −80 °C. Genomic DNA from these pellets was extracted with the DNeasy Blood & Tissue Kit (Qiagen). Amplification of the sgRNA barcodes and indexing of each sample was performed via two-step PCR as previously described (*20, 80*). To determine differences in sgRNA composition between samples, deep sequencing was performed using the Illumina Nextseq platform (single-ended 75 bp). As previously described (*20, 80*), barcoded reads were converted to guide-level counts and the fractional representation of each sgRNA construct was found by dividing the count of each sgRNA in a sample by the sum of all sgRNA counts in that sample. Construct-level depletion scores were collapsed to gene-level depletion scores by taking the average depletion score across five sgRNA constructs. All depletion/enrichment effects were reported as log2 ratios.

### Resistance potential

MIA PaCa-2 cells were plated in six-well plates at a density of 500 cells per well. The following day, cells were treated with increasing concentrations of SCH772984, with all conditions performed in technical triplicate. Following two weeks of treatment, plates were rinsed with PBS and fixed and stained with 0.5% (wt/vol) crystal violet in 6.0% (vol/vol) glutaraldehyde solution (Thermo Fisher Scientifics) for 30 min at room temperature. Plates were rinsed in distilled H_2_O and photographed the following day. Colonies were hand-counted using scanned images at 400x, and resistance potential was defined as the number of colonies in drug-treated conditions relative to DMSO-treated cell.

### Long-term tumoroid assay

Colorectal cancer tumoroids, generated and maintained as previously described (*61*), we treated with DMSO alone (1:1,000), Senexin A (1 μM), SCH772084 (1 μM), or a combination of both SCH772084 (1 μM) and Senexin A (1 μM). As previously described (*61*), tumoroids were passaged for up to eight weeks, with counts performed at weekly time points.

## Funding

National Institutes of Health F32CA180569 (DPN)

National Institutes of Health R01CA207083 (KCW)

National Institutes of Health R01CA263593 (KCW)

National Institutes of Health R01CA175747 (CJD)

National Institutes of Health U01CA199235 (CJD)

National Institutes of Health P01CA203657 (CJD)

National Institutes of Health R35CA232113 (CJD)

Department of Defense Lung Cancer Research Program W81XWH2110362 (KCW)

Duke University School of Medicine/Duke Cancer Institute start-up funds (KCW)

American Cancer Society PF 18-061 (AMW)

## Author contributions

Conceptualization: DPN, CAM, EFP, JJS, TER, CJD, DJT, and KCW

Methodology: DPN, CAM, EFP, JJS, TER, CJD, DJT, and KCW

Investigation: CAM, AMW, AB, JCR, CGC-S, AES, CW, MC, CBW, DEK, and MP

Visualization: DPN

Supervision: KCW, EFP, JJS, TER, CJD, DJT

Writing—original draft: DPN, KCW

Writing—review & editing: All authors

## Data and materials availability

All data needed to evaluate the conclusions in the paper are present in the paper and/or the Supplementary Materials. Request for materials can be provided by KCW pending scientific review and a completed material transfer agreement. Requests for the data should be submitted to.

