## Supplemental Figures for "Mediator Kinase Inhibition Impedes Transcriptional Plasticity and Prevents Resistance to ERK/MAPK-Targeted Therapy in *KRAS*-Mutant Cancers"

**This PDF file includes:**

Figs. S1 to S5

**Other Supplementary Materials for this manuscript include the following:**

Tables S1 to S10

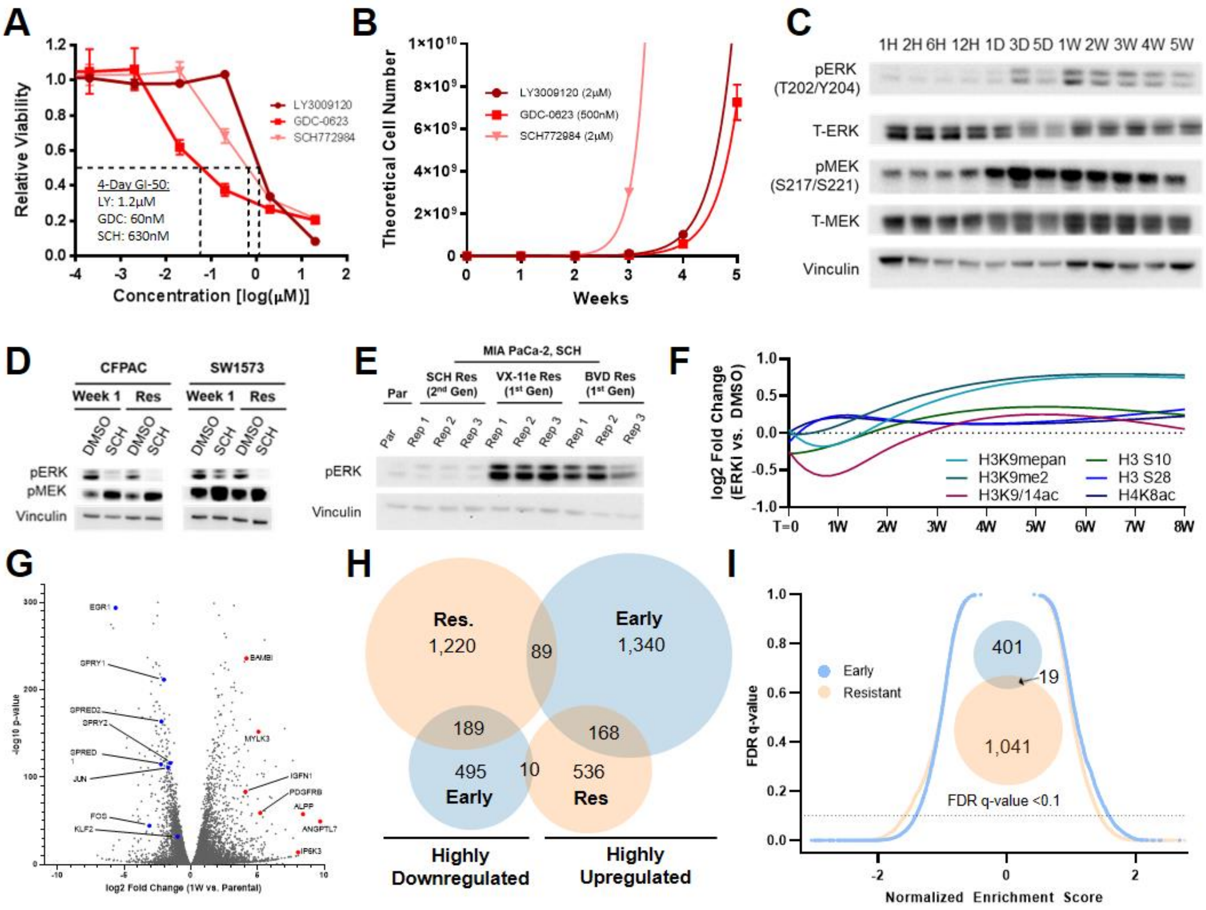


**Fig. S1. Transcriptional and epigenetic changes associated with ERK/MAPK resistance, related to Figure 1. (A)** Eight-point growth inhibition assay of MIA PaCa-2 cells treated with the indicated MAPK inhibitors at increasing concentrations, each condition performed in triplicate. **(B)** TTP assay of MIA PaCa-2 cells treated with the indicated MAPK inhibitors for up to five weeks, each condition performed in triplicate. **(C)** Immunoblot of MIA PaCa-2 cells treated with SCH772984 (1 μM) for the indicated duration. **(D)** Immunoblot of CFPAC-1 and SW1573 cells treated with either DMSO (1:1,000) or SCH772984 (1 μM) for one week or eight weeks (stable resistance). **(E)** Immunoblot of treatment-naïve parental and evolved resistant MIA Paca-2 cells. Evolved resistant cells were treated for eight weeks with SCH772984 (1 μM), VX-11e (1 μM), or BVD-523 (1 μM), respectively, with each condition performed in triplicate. **(F)** Restricted cubic splines of protein expression changes of various histone markers from RPPA. **(G)** Volcano plot indicating gene expression changes of MIA PaCa-2 cells treated with SCH772984 (1 μM) for one week (early response) relative to DMSO (1:1,000). **(H)** Venn diagram indicating highly upregulated or downregulagated transcripts (|Log2FC| >1.5, p<0.001) after treatment with SCH772984 (1 μM) for one week (early) or eight weeks (stable resistance) relative to DMSO (1:1,000). **(I)** Reverse volcano plot depicting gene set enrichment analysis using gene sets from the MsigDB Biologic Process Ontology based on the gene expression data MIA PaCa-2 cells treated with SCH772984 (1 μM) for one week or eight weeks relative to DMSO (1:1,000); Venn diagram depicting enriched gene sets with an FDR <0.1 (insert).

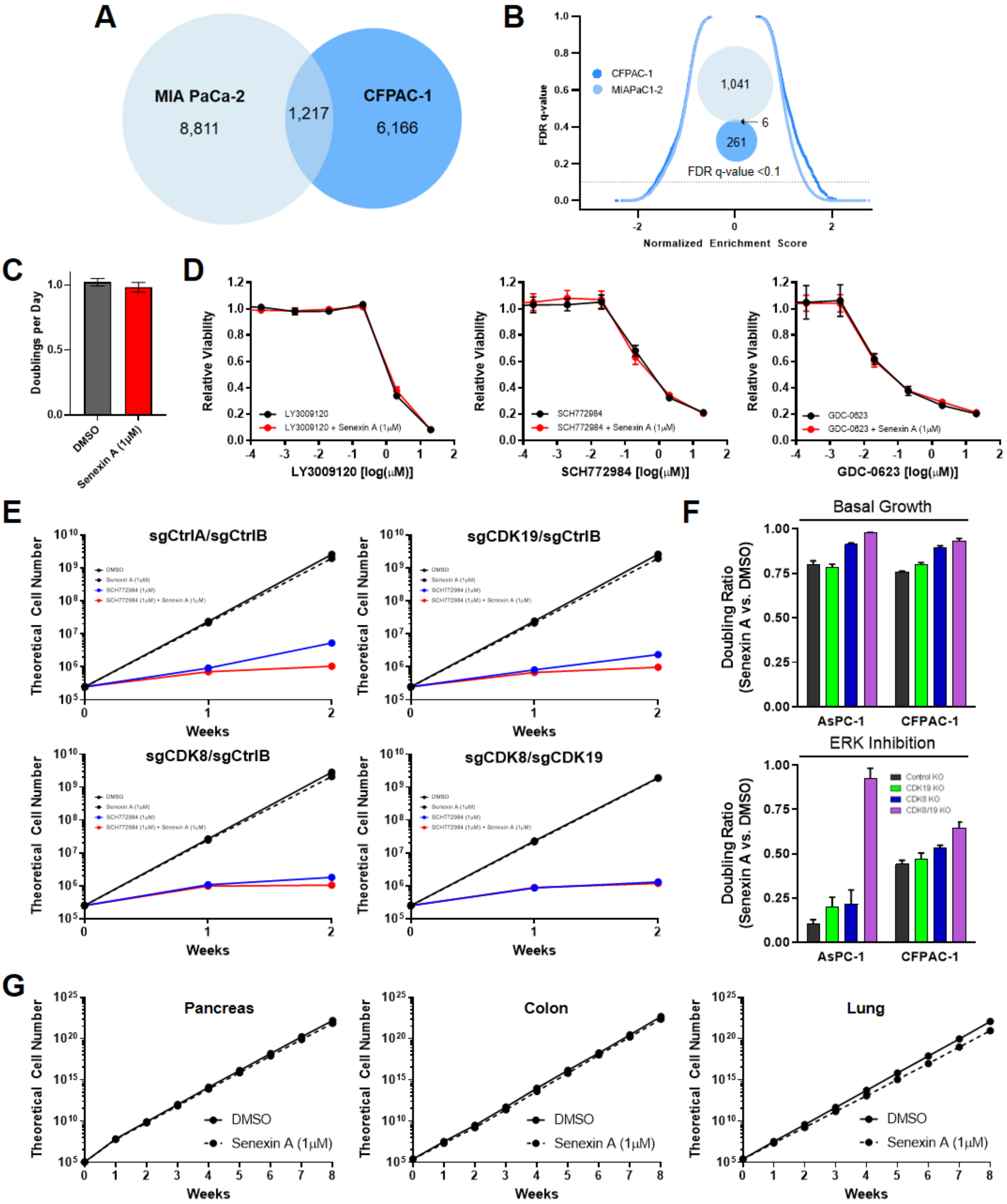


**Fig. S2. Mediator kinase inhibition prevents long-term adaptations to ERK/MAPK inhibition, related to Figure 2. (A)** Venn diagram indicating differentially expressed genes ($p < 10^{-3}$) in MIA PaCa-2 cells and CFPAC-1 cells treated with SCH772984 (1 µM) for eight weeks (stable resistance) compared to DMSO (1:1,000), each condition performed in triplicate. **(B)** Reverse volcano plot depicting gene set enrichment analysis using gene sets from the MsigDB Biologic Process Ontology based on the gene expression data in Figure S2A; Venn diagram depicting enriched gene sets with an FDR <0.1 (insert). **(C)** Mean cell doublings per day of MIA PaCA-2 cells treated with either Senexin A (1 µM) or DMSO (1:1,000) for three weeks, each condition performed in triplicate. **(D)** Eight-point growth inhibition assay of MIA PaCa-2 cells treated with the indicated MAPK inhibitors at increasing concentrations in the background of Senexin A (1 µM) or DMSO (1:1,000), performed in triplicate. **(E)** Short-term (two week) TTP assay of MIA-PaCA-2 cells with the indicated genetic perturbations treated according to the indicated condition, with each condition performed in triplicate. **(F)** Doubling ratio of AsPC-1 cells and CFPAC-1 cells over the short-term TTP assay based on the indicated genetic perturbations and treated according to the indicated conditions, demonstrating the effect of CDK8/19 inhibition either in basal growth conditions (top) or in the presence of SCH772984 (1 µM; bottom), each condition performed in triplicate. **(G)** TTP assay of MIA PaCa-2, SW620, and SW157 cells treated with DMSO (1:1,000) or Senexin A (1 µM) for eight weeks, each condition performed in triplicate.

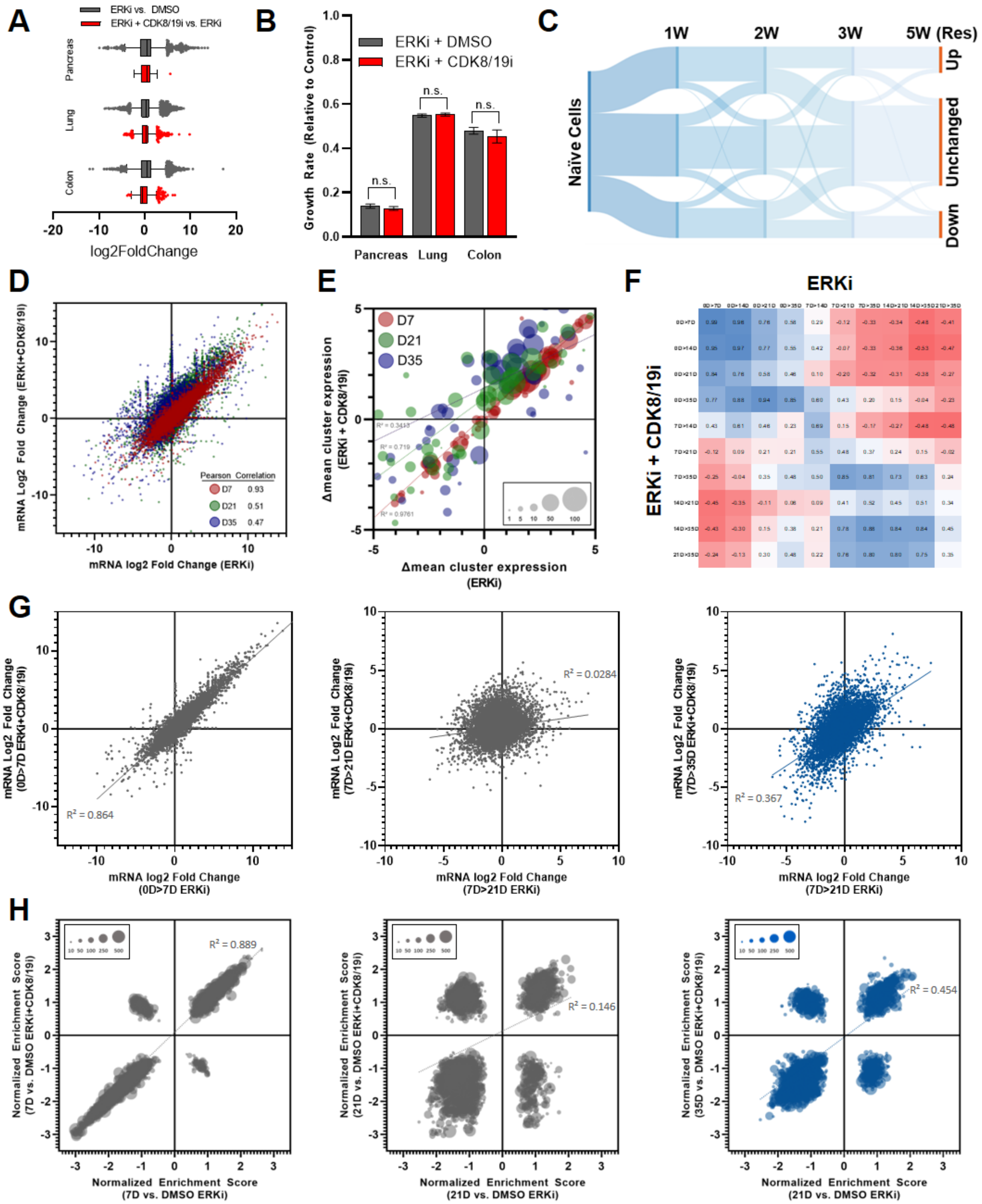


**Fig. S3. Mediator kinase inhibition prevents the downstream transcriptional adaptations necessary for resistance to ERK/MAPK inhibition, related to Figure 3.** A) Violin plot depicting transcriptome-wide significant (p <0.001) gene expression changes in MIA PaCa-2 cells, SW1753 cells, and SW620 cells treated with SCH772984 (1 µM) in combination with DMSO (1:1,000) versus Senexin A (1 µM). Statistical significance determined by Wald test using the Benjamini and Hochberg method to correct for multiple hypothesis testing. **(B)** Growth rate of MIA PaCa-2 cells, SW1753 cells, and SW620 cells treated with SCH772984 (1 µM) in combination with DMSO (1:1,000) versus Senexin A (1 µM). **(C)** Sankey plot demonstrating the magnitude and directionality of gene expression changes ($p < 0.001$) of MIA PaCa-2 cells treated with SCH772984 (1 µM) compared to DMSO (1:1,000) over a five-week time course. **(D)** Scatter plot comparing gene-level expression changes of MIA PaCa-2 cells treated according to the indicated conditions at the indicated time points. **(E)** Bubble charts reflecting the mean expression changes of each cluster in the indicated treatment conditions at the indicated time points; the size of each bubble reflecting the number of genes in each cluster. **(F)** Heatmap demonstrating the Pearson correlation of gene-level expression changes in MIA PaCa-2 cells at all time point comparisons along a five-week time course according to the indicated treatment conditions. **(G)** Scatter plots comparing gene-level expression changes of MIA PaCa-2 cells treated according to the indicated conditions at various indicated time points along a five-week time course. **(H)** Bubble charts reflecting the normalized enrichment score of all gene sets from the MsigDB Biologic Process Ontology generated from gene expression changes of MIA PaCa-2 cells treated in the indicated conditions at various indicated time points along a five-week time course. For all subfigures, each condition was performed in technical triplicate.

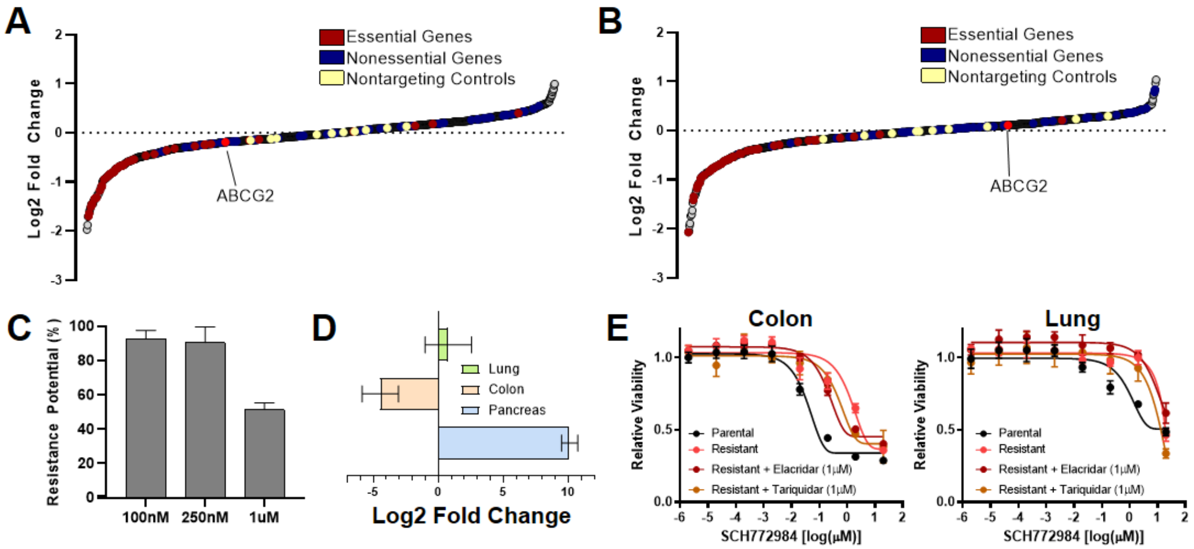


**Fig. S4. Resistance to ERK/MAPK inhibition is characterized by transcriptional plasticity, related to Figure 4. (A, B)** Gene-level representation of essential phenotypes in evolved resistant MIA PaCa-2 cells (A) and treatment-naïve parental cells (B) treated with DMSO (1:1,000) at the screen midpoint (19 days), ranked by their mean log2-transformed gene score across duplicates. **(C)** Potential of individual MIA PaCa-2 cells to develop resistance to increasing concentrations of SCH772984. **(D)** ABCG2 mRNA expression change of MIA PaCa-2 cells, SW1573 cells, and SW620 cells treated with SCH772984 (1 μM) for eight weeks (terminally resistant) relative to their treatment-naïve, parental counterparts. **(E)** Eight-point growth inhibition assay of parental and evolved resistant SW620 cells (left) and SW1573 cells (right) treated with increasing concentrations of SCH772984 alone or in the background of Elacridar (1 μM) or Tariquidar (1 μM).

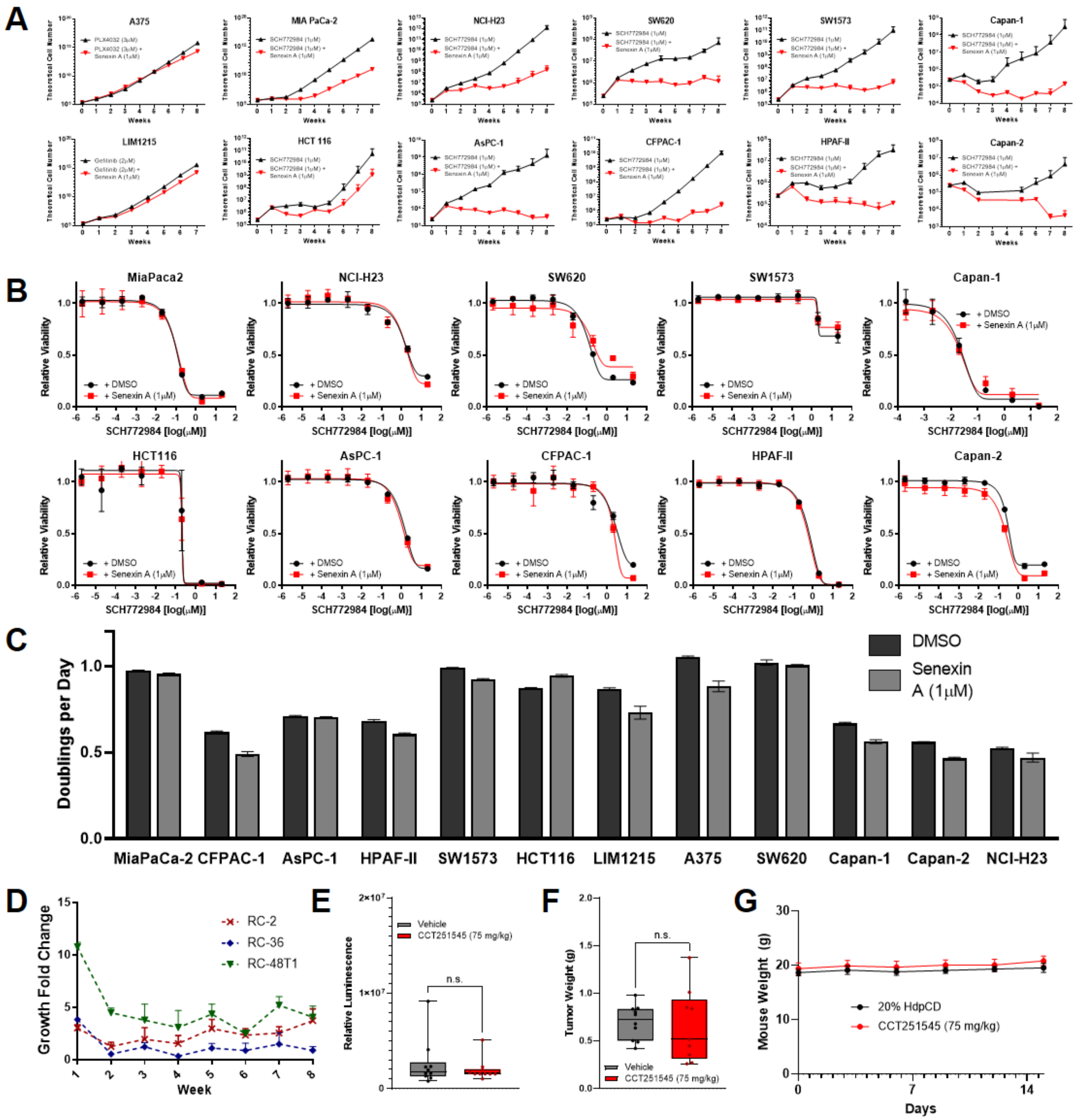


**Fig. S5. Translational models of combined ERK and Mediator kinase inhibition, related to Figure 5.** (A) TTP assays for 12 indicated cell lines (10 *KRAS*-mutant and two *KRAS* wild-type) treated with SCH772984 (1 µM) alone or in combination with Senexin A (1 µM). **(B)** Eight-point growth inhibition assay of 10 *KRAS*-mutant cell lines treated with increasing concentrations of SCH772984 (1 µM) in the background of DMSO (1:1,000) versus Senexin A (1 µM). **(C)** Mean doublings per day (with error bars reflecting SD) of 12 indicated cell lines treated with DMSO (1:1,000) alone or Senexin A (1 µM) alone over the eight-week TTP assay. **(D)** TTP assays for rectal cancer tumoroids treated with SCH772984 (100nM) combined with Senexin A (1 µM) for the entire eight-week TTP assay. **(E, F)** Box plots demonstrating whole body luminescence **(E)** and tumor weights **(G)** after orthotopic implantation of $10^3$ $2.1.1^{syn\_Luc}$ cells into FVB/n mice and treatment with vehicle (20% HpBCD) or CCT251545 (75 mg/kg) for three weeks, 10 mice per group. **(G)** Mean mouse weights throughout treatment described in Figure 5E, F.
